# Gene flow improves fitness at a range edge under climate change

**DOI:** 10.1101/399469

**Authors:** Megan Bontrager, Amy L. Angert

## Abstract

Populations at the margins of a species’ geographic range are often thought to be poorly adapted to their environment. According to theoretical predictions, gene flow can inhibit these range edge populations if it disrupts adaptation to local conditions. Alternatively, if range edge populations are small or isolated, gene flow can provide beneficial genetic variation, and may facilitate adaptation to environmental change. We tested these competing predictions in the annual wildflower *Clarkia pulchella* using greenhouse crosses to simulate gene flow from sources across the geographic range into two populations at the northern range margin. We planted these between-population hybrids in common gardens at the range edge, and evaluated how genetic differentiation and climatic differences between edge populations and gene flow sources affected lifetime fitness. During an anomalously warm study year, gene flow from populations occupying historically warm sites improved fitness at the range edge, and plants with one or both parents from warm populations performed best. The effects of the temperature provenance of gene flow sources were most apparent at early life history stages, but precipitation provenance also affected reproduction. We also found benefits of gene flow that were independent of climate: after climate was controlled for, plants with parents from different populations performed better at later lifestages than those with parents from the same population, indicating that gene flow may improve fitness via relieving homozygosity. Further supporting this result, we found that increasing genetic differentiation of parental populations had positive effects on fitness of hybrid seeds. Gene flow from warmer populations, when it occurs, is likely to contribute adaptive genetic variation to populations at the northern range edge as the climate warms. On heterogeneous landscapes, climate of origin may be a better predictor of gene flow effects than geographic proximity.

**Impact summary:** What limits species’ geographic ranges on the landscape? One process of interest when trying to answer this question is gene flow, which is the movement of genetic material between populations, as might occur in plants when seeds or pollen move across the landscape. One hypothesis that has been proposed is that gene flow from populations in other environments prevents populations at range edges from adapting to their local habitats. Alternatively, it has been suggested that these populations might benefit from gene flow, as it would provide more genetic material for natural selection to act upon.

We tested these predictions in an annual wildflower, *Clarkia pulchella*. We simulated gene flow by pollinating plants from the range edge with pollen from other populations. Then we planted the resulting seeds into common gardens in the home sites of the range edge populations and recorded their germination, survival, and reproduction. The weather during our experiment was much warmer than historic averages in our garden sites, and perhaps because of this, we found that gene flow from warm locations improved the performance of range edge populations. This result highlights the potential role of gene flow and dispersal in aiding adaptation to warming climates. We also found some positive effects of gene flow that were independent of climate. Even after we statistically controlled for adaptation to temperature and precipitation, plants that were the result of gene flow pollinations produced more seeds and fruits than plants with both parents from the same population. Rather than preventing adaptation, in our experiment, gene flow generally had positive effects on fitness.

## Introduction

Species are limited in their geographic extents on the landscape. In many cases, the limits of species’ geographic distributions are the result of niche limitation, rather than simply an inability to disperse to suitable areas beyond their current distribution (Lee-Yaw et al., 2016). This raises the question of what prevents populations on the range periphery from adapting to sites beyond the range edge (Antonovics, 1976; Bridle and Vines, 2007), particularly when boundaries are not co-incident with an abrupt shift in the abiotic environment. The putative causes of limits to adaptation at the range edge hinge upon demographic and genetic features of metapopulations (Sexton et al., 2009).

If range limits represent limits to adaptation, this could be the result of insufficient genetic variation in range edge populations. Limited genetic variation at range edges is predicted because range edge populations are often characterized as small, with more frequent or severe changes in population size and elevated rates of turnover relative to central populations (Vucetich and Waite, 2003). Populations that are on the leading edge of range expansions may also exhibit patterns of low genetic variation as a result of successive founder events (Pujol and Pannell, 2008). Significant declines in neutral genetic variation near range edges is a common (though not ubiquitous) pattern (Eckert et al., 2008; Pironon et al., 2017), indicating that some of these processes are likely to affect some range edges in some species. If the observed declines in neutral variation also reflect reduced adaptive genetic variation, this might result in marginal populations being less locally adapted when compared to central populations, as they have less capacity to respond to local selection pressures. Maladaptation is expected to lead to poor demographic performance, reducing colonization opportunities in sites beyond the range, and potentially creating (or reinforcing) a range edge at equilibrium along an environmental gradient (Kirkpatrick and Barton, 1997).

Swamping gene flow is another often-invoked hypothesis for how equilibrial range limits might form and persist (Lenormand, 2002; Sexton et al., 2009). Under swamping gene flow, peripheral populations are unable to adapt to their local conditions because they experience maladaptive gene flow from central populations (Kirkpatrick and Barton, 1997). This process is predicted to occur when populations are arranged along an environmental gradient where individuals are well-adapted and abundant in the center of that gradient. Because of this asymmetry in abundance, net gene flow is asymmetric and brings alleles that are adaptive in central environments to edge populations, disrupting local adaptation to edge environments. This causes edge populations to become demographic sinks, where death rates exceed birth rates, and prevents further range expansion. According to this model, the fitness of edge populations will depend upon the rate of gene flow from center to edge as well as the steepness of the environmental gradient (i.e., the magnitude of environmental differences between the sources of the gene flow and the recipient populations).

Comprehensive empirical tests of the swamping gene flow hypothesis are difficult to conduct because they require demonstrating both the negative effects of gene flow on edge populations as well as the occurrence of asymmetric gene flow on the landscape. Evidence to-date indicates that in some systems swamping gene flow might limit adaptation along geographic gradients (Paul et al., 2011) or between habitat types (Anderson and Geber, 2010; Farkas et al., 2016) and may sometimes limit the geographic range (Fedorka et al., 2012; Holliday et al., 2012). However, in other systems there are no detectable fitness costs of gene flow across environmental gradients (Emery, 2009; Moore and Hendry, 2009; Samis et al., 2016) and strong local adaptation persists despite gene flow (Yeaman and Jarvis, 2006; Gould et al., 2014). Outbreeding depression may generate patterns similar to those resulting from the disruption of local adaptation, but the effects of genetic incompatibilities can be discerned from those of swamping by experimental designs that allow for decoupling of environmental and genetic differentiation.

Most theory about swamping gene flow at range edges has been developed with the assumption of smooth environmental gradients underlying the range, however, this assumption is unrealistic for most species. Topography, continentality, and other landscape features make transects from range centers to edges heterogenous with regards to climate. Other habitat variables, such as soil type or the biotic community (which may mediate responses to climate in addition to imposing selection on their own), are also likely to be spatially heterogeneous. This complicates predictions of the swamping gene flow hypothesis: range edge populations may experience gene flow from environmentally divergent neighboring populations, or environmentally similar central populations, as well as combinations falling anywhere in between. In this case, geography cannot be used as a proxy for predictions about the effects of gene flow, rather, these predictions must be informed by the environmental differences between populations. Gene flow between populations in similar environments may be beneficial, even when populations are geographically disparate, because gene flow can allow for the spread of environment-specific beneficial alleles that arise in a single population (Sexton et al., 2011). Abundant-center distribution patterns and asymmetric gene flow have been documented in some species, but are not ubiquitous (Sagarin and Gaines, 2002), perhaps at least in part as a result of complex environmental gradients.

In addition to contributing alleles that are adaptive or maladaptive in a given environment, gene flow may provide relief from homozygosity caused by drift or inbreeding. Gene flow is expected to increase heterozygosity and reintroduce variation that can allow for masking or purging of fixed deleterious alleles. As a result, gene flow can improve fitness in peripheral populations (Sexton et al., 2011). The extent to which gene flow causes heterosis depends upon the genetic divergence of populations (Ingvarsson and Whitlock, 2000), but not explicitly on the magnitude of the environmental differences between the source and recipient of gene flow, though environmental differences are correlated with genetic differentiation in some species (Sexton et al., 2014).

Gene flow may also be beneficial when maladaptation arises due to disequilibrium between a populations’ optimal conditions and the environment. This could occur when a species is undergoing a range expansion, or when the environmental landscape is moving out from under individuals, as is occurring under climate change (Aitken and Whitlock, 2013). If a population is locally adapted to historic conditions in a site, and the environment changes rapidly, then gene flow from populations with historic conditions that are more similar to these new local conditions is expected to improve population performance.

To investigate how gene flow affects peripheral populations, we simulated gene flow among populations spanning the northern half of the range of an annual wildflower, *Clarkia pulchella*, and measured lifetime fitness of individuals in two common gardens at the species’ northern range edge. We asked 1) Are range edge populations of *C. pulchella* locally adapted? 2) What climatic factors predict fitness at the northern range edge? 3) Does gene flow positively or negatively affect edge populations? and 4) How does the effect of gene flow from other populations depend upon the genetic differentiation and climatic distances of these populations? Under conditions where the range edge is not at equilibrium with climate, we expect that gene flow from sites that are historically similar to the experimental conditions will improve performance. Under conditions in which this species’ range is at equilibrium with climate, and if this edge is limited by adaptation, we expect that gene flow from populations in similar climates will have a positive effect on fitness via heterosis or the contribution of adaptive alleles, but that gene flow from strongly contrasting climates will be detrimental. If populations have genetic incompatibilities (which need not be the result of divergent selection, but could simply be the result of drift under prolonged separation) then we expect the offspring of crosses between populations that are more genetically divergent to perform worse, regardless of the conditions of the test environment. However, if homozygosity is negatively related to fitness we expect a greater benefit from gene flow between more genetically divergent populations.

## Methods

### Study system, seed collection, and site selection

*Clarkia pulchella* Pursh (Onagraceae) is a winter annual that grows on sparsely vegetated, south-facing slopes with low canopy cover throughout eastern Washington and Oregon, Idaho, and western Montana (United States) and southeastern British Columbia (Canada). This species germinates in fall and overwinters as a seedling before flowering in late May, June, and early July. It has no observed seed dormancy, but seeds will not germinate immediately upon dehiscing and require an after-ripening period of several weeks. It has showy pink flowers and is visited by a diverse array of pollinators though it has some capacity to self-pollinate in the absence of pollinators or mates (MacSwain et al., 1973; Bontrager et al., 2018). Individual plants that survive to reproduction typically produce 1–35 flowers (mean = 3.11, sd = 3.07; M. Bontrager, unpublished data).

Seeds of *C. pulchella* were collected from 15 populations in July 2014 (Figure 1A; Table S1). We used the two northwestern-most populations of the continuous distribution of *C. pulchella* as common garden sites (hereafter referred to as focal populations). Other populations (hereafter, donor populations) were selected with the goal of sampling representative variation in major climatic axes (temperature, precipitation, and seasonality of these variables; Figure 1B) across the northern half of the species’ range. In each of these populations, seeds were collected haphazardly from plants spaced >0.5 m apart.

**Figure 1.**
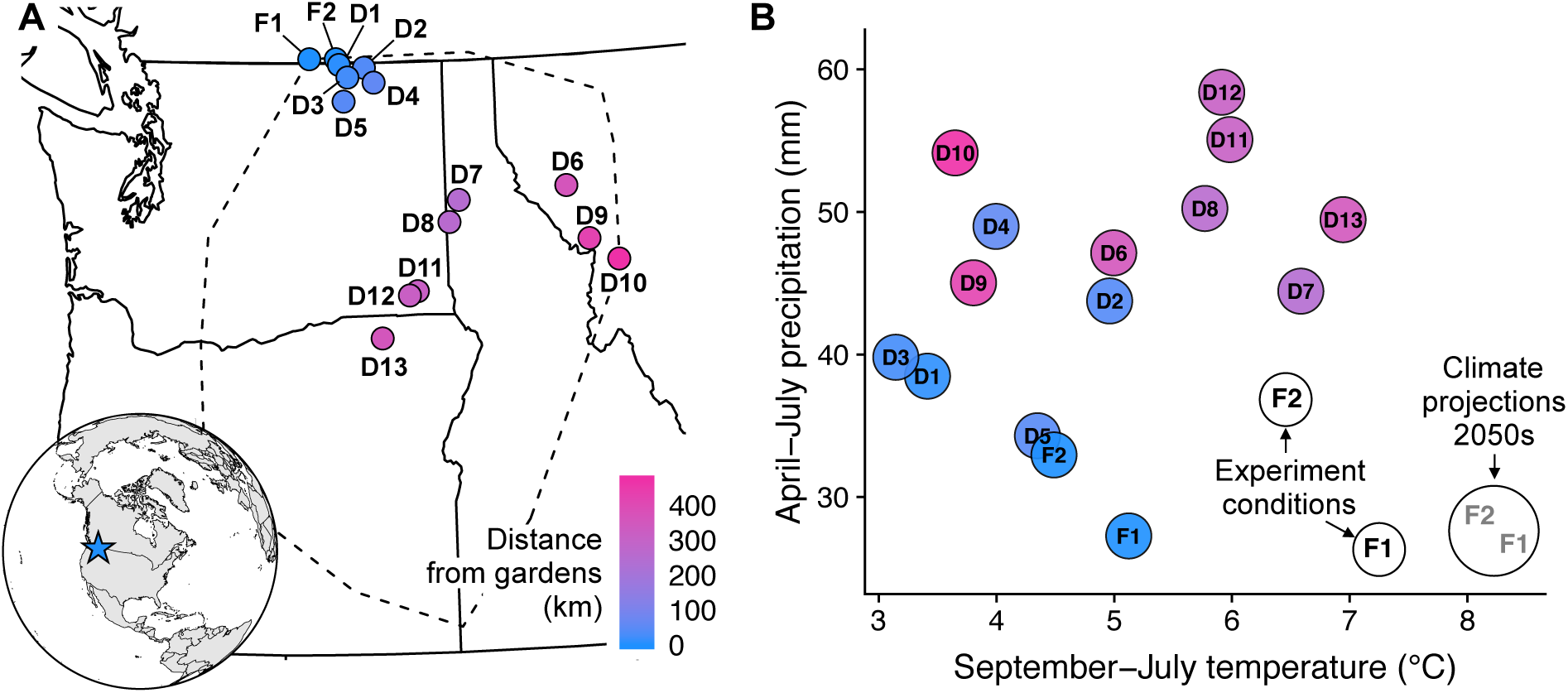
Geographic locations and climate averages of populations used in this experiment. **(A)** Populations span the northern half of the geographic range of *Clarkia pulchella* (indicated by the dashed line; the range is represented by the extent of localities in the Consortium of Pacific Northwest Herbaria database). Focal population sites (where common gardens were installed) are indicated by “F” and donor population sites are indicated by “D”. Identifying codes for each population correspond to the Map ID column in Table S1. **(B)** Temperature and precipitation conditions of common gardens and source provenances (calculated from monthly PRISM data; PRISM Climate Group 2017). Colored circles represent the 1951-1980 average September-July temperature and summed April-July precipitation in the home site of each population. Open circles represent weather conditions during the experiment. Focal populations are historically intermediate relative to donor populations in average historic temperature (x-axis), but are the from the driest sites of any population used in the experiment (y-axis). Conditions in common gardens during the experiment were hot relative to normal conditions at those sites, and hot and dry relative to average conditions of all populations in the experiment. Conditions during the experiment were not quite as warm as projected temperatures for 2055 under the HadGEM2-ES RCP 4.5 model (grey text, downscaled using ClimateNA, Wang et al. 2016).

### Greenhouse generation and crossing design

We grew field-collected seeds in the greenhouse and implemented a controlled crossing design to generate seeds for the field transplant. We planted maternal families from each population into 22 randomized blocks, with each block containing one family from each population. Two types of crosses were performed: “within-population” crosses and “between-population” crosses. For within-population crosses, dams were pollinated using pollen from the plant of the same population in the subsequent block (Figure 2A). Each plant from each of the 15 populations was therefore used as both a sire and a dam with other plants from the same population. For between-population crosses, flowers on plants from the two focal populations within each block were pollinated using each of the donor plants in that block (Figure 2B). These crosses simulate an early stage of gene flow: the progeny of a mating event that is the result of long distance pollen dispersal (or the progeny of a cross between a native individual and a recent immigrant). We collected the seeds from crosses as fruits ripened. Further details about our greenhouse conditions and crossing design are in Supplementary Methods 1.

**Figure 2.**
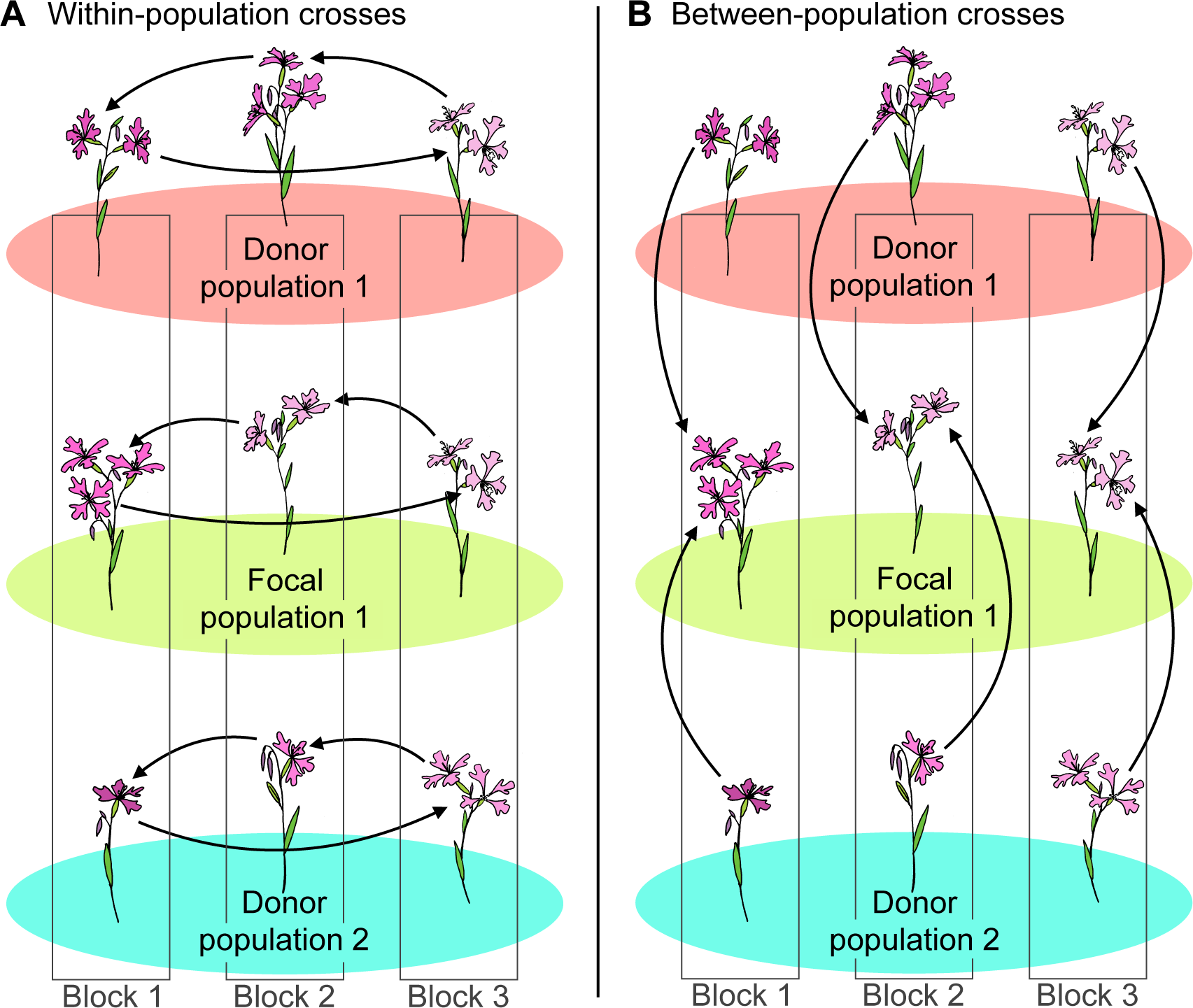
Schematic of the greenhouse crossing design to generate seeds for our common gardens. Three blocks are diagrammed here, but crosses were performed in 22 blocks and seeds from 15 blocks were transplanted into the field. **(A)** Within-population crosses: for each seed family from each of 15 populations (three are shown), plants were crossed in a “daisy chain” design, in which each plant was hand-pollinated using pollen from another individual of the same population from the subsequent block. **(B)** Between-population crosses: we used pollen from each of 13 donor populations (two are shown) to pollinate flowers on plants from each of the two focal populations (one is shown). Each focal plant served as a dam for multiple between-population crosses, that is, each focal seed family had one flower pollinated by a plant from each of 13 donor populations and from the other focal population.

### Common garden design and installation

For the transplant, we used seeds from 15 of our greenhouse blocks, and substituted seeds from the same type of cross from other greenhouse blocks when they were unavailable from our primary 15. Seeds were glued to toothpicks to expedite planting and monitoring in the field. Two seeds were glued to each toothpick with a tiny dab of water-soluble glue (when seeds were limited, just one seed was glued to each toothpick). At each of the two sites, toothpicks were planted into 10 fully randomized plots. Each plot contained two toothpicks from each cross type from each of the 15 replicates. We only planted between-population crosses with local dams at each of the two focal sites (i.e., Blue Lake plots only contained between-population crosses performed on Blue Lake plants, and Rock Creek plots only contained between-population crosses performed on Rock Creek plants). Within-population crosses from all populations were planted out at both sites. Therefore, each plot contained two replicates of each of 15 crosses from 29 cross types (14 between-population groups and 15 within-population groups). For some cross types, less than 15 families had sufficient seeds for the full design, therefore each plot contained 832 toothpicks at Rock Creek and 836 toothpicks at Blue Lake. In total, our design included 16,680 toothpicks and 32,755 seeds. Seeds were planted in September 2015 and watered once in late October, though at that time most seeds that were checked had already begun germinating. Plots were protected with wire cages; further details of transplant installation are in Supplementary Methods 2.

### Monitoring and measuring

Germination was censused 16-20 November 2015. We documented the emergence of either 0, 1, or 2 germinants at each toothpick. If two germinants were present, these were randomly thinned so that just one remained. Overwinter survival was assessed 17-21 March 2016. During both germination and overwinter survival censuses, plant size was measured as the distance in mm from the tip of one leaf to the tip of the other (most plants had just one pair of leaves). Further details of monitoring and measuring are provided in Supplementary Methods 3. In early June 2016 we censused the spring survival of all plants and began measuring reproduction every 2-3 days. As a proxy for seed production, we measured the ovary length of each flower produced on each plant. Pollinators were excluded from our plots to prevent the escape of non-native genotypes, so we calibrated a conversion from ovary length to seed production using hand pollinations (linear regression of seed production on ovary length: 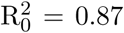, p < 0.0001; Supplementary Methods 4). We continued reproductive censuses until all plants had senesced, but reduced census frequency when flowering slowed in July.

### Climatic and genetic differentiation

We compiled monthly temperature and precipitation data from 1951-1980 for all seed sources, as well as the gardens during the months of the experiment (September 2015-July 2016) from PRISM (PRISM Climate Group, 2017). We calculated historic (pre-warming, 1951-1980) climate averages for each site, which we compared to conditions experienced by plants during the transplant in our analyses. Our decision to use normals from this time window, rather than a more recent time window, reflects an assumption that populations have been limited in their capacity to adapt to changing conditions, such that historic normals best represent each population’s climate optima. However, we note that contemporary normals (1984-2013) are highly correlated with those that we used (September-July temperature: r = 0.99, p < 0.0001; April-July precipitation: r = 0.96, p < 0.0001, Figure S1, S2), so our choice of time period is unlikely to affect our results. Inter-annual climate variability was very similar in each of the seed collection sites (data not shown) so we do not further consider variability in climate, and focus on averages only. Pairwise population genetic differentiation (F_ST_) was calculated from 2982 SNPs that were genotyped in up to 12 parental individuals from each population (genotyping and alignment methods are described in Bontrager and Angert 2018). F_ST_ was calculated using the implementation of Weir and Cockerham (1984) in the R package hierfstat (Goudet and Jombart, 2015).

### Statistical analyses

#### Did local populations outperform foreign populations?

We tested whether local populations were, on average, superior to foreign populations using within-population crosses only. We compared lifetime fitness of local vs. foreign individuals in a generalized linear mixed model (GLMM) with a zero-inflated negative binomial distribution using the package glmmTMB (Brooks et al., 2017). These zero-inflated models allow specification of fixed effects for both the zero-inflation part of the model (the probability of a non-zero value) as well as the conditional part of the model (the effect on the response after accounting for zero-inflation). Generally, we consider the zero-inflation part of the model to reflect early lifestages, as the majority of plants that produced zero seeds did so as a result of failing to germinate or survive winter. The conditional part of the model may reflect both differences during reproduction as well as differences among individuals accumulated across all lifestages. In addition to testing for local adaptation represented by lifetime fitness, we tested whether local populations performed better than foreign populations at any component lifestage: germination, size after germination, overwinter survival, size after winter, fruit count, and estimated seed production. Plant size was modeled with a Gaussian response distribution, germination and survival with binomial response distributions and logit link functions, and fruit counts and seed production with zero-inflated negative binomial response distributions and log link functions. For component lifestage analyses, we included only individuals that had survived to the preceding census, and always included plant size at the previous census to account for differences that had accumulated at earlier lifestages. For all of these models we initially included a random effect structure of block within site, dam within dam population, and sire within sire population. However, models of later lifestages and lifetime fitness frequently failed to converge with this parameterization. When convergence failed, we reduced random effects to only block within site and sire population. Predictions and confidence intervals were visualized with ggeffects (Lüdecke, 2018).

#### Does climate of origin explain performance in common gardens?

We built GLMMs using the methods described above to evaluate the effects of mismatch between the experiment conditions and population’s historic climates on lifetime fitness and all fitness components, again using only within-population crosses. We calculated the absolute difference between the garden conditions and the historic temperature and precipitation of each source population. We use absolute differences because we expect that mismatch in either direction along a climate axis will negatively impact fitness. Very few source populations were from sites drier or hotter than conditions during the experiment, so absolute differences mostly result from source populations being historically cooler or wetter than the experiment. Our lifetime fitness model included absolute differences in temperature (for the experiment duration, September-July) in both the conditional and zero-inflation parts of the model, as well as absolute differences in spring and summer precipitation (April-July) in the conditional part of the model only. We isolated the specific lifestages affected by each of these climatic predictors using the methods described above. In lifestage-specific analyses, we calculated climate differences using only the months in each census window. In all analyses, all continuous predictors were scaled and centered.

#### Does gene flow help or hurt edge populations?

Based on the results of the analyses above, we expected that gene flow from some populations was likely to confer benefits by contributing adaptive genetic variation to focal populations experiencing an anomalous climate. To evaluate whether there were benefits of gene flow that were independent of these climate effects, we calculated the midparent historic temperature average for all individuals (that is, the average temperature of dam and sire sites) and then calculated the absolute difference between this temperature and the experimental temperature (Figure S3). We calculated a metric of absolute midparent precipitation difference (summed over April-July) in the same manner. We used GLMMs as described above to test for an effect of gene flow in addition to an anticipated effect of midparent climate differences on lifetime fitness and each component lifestage. In these models, gene flow was included as a categorical fixed effect (within-population cross vs. between-population cross) along with midparent temperature and precipitation differences. We included gene flow and temperature differences in both the conditional and zero-inflation parts of the lifetime fitness model, and precipitation differences in only the conditional part. We had difficulty disentangling effects of precipitation differences and gene flow (see Results), so we ran these models with and without precipitation differences.

#### Do the effects of gene flow depend upon the genetic differentiation between focal and donor populations?

We examined whether the genetic differentiation (F_ST_) between the two parental populations of the between-population crosses positively or negatively affected offspring fitness. We could only estimate genetic differentiation between parental populations for individuals with parents from different populations, so in these analyses we use between-population crosses only. We built zero-inflated GLMMs as described above using lifetime fitness and included predictors of absolute midparent temperature and precipitation differences as well as F_ST_. We also tested the effects of these parameters on each component lifestage. Our ability to detect significant effects of climate in full models was limited, likely due to the narrow range of midparent climatic variability across between-population crosses, so we also built separate models of each of our three predictors on lifetime fitness and each lifestage. All statistical analyses were implemented in R version 3.4.3 (R Core Team, 2017).

## Results

### Climate of origin explains performance in common gardens

Local populations were not superior to the average foreign population in their cumulative fitness across all lifestages, or in any component lifestage (Figure S4, Table S2). Populations that were best matched to experimental temperatures performed best in our gardens; lifetime fitness declined with increasing absolute temperature differences between the source and the experimental conditions (Figure 3A). This occurred via effects on both the probability of producing any seeds (the zero-inflation part of the model; *β* = −0.341, SE = 0.033, *P* < 0.001; Figure 3B), and the number of seeds produced (the conditional part of the model; *β* = −0.114, SE = 0.050, *P* = 0.022; Figure 3C). Local populations, which are historically intermediate in temperature (Figure 1B), were mismatched from the experiment conditions and performed worse than populations from warmer sites that were more climatically similar to the garden conditions (Supplementary Analysis 1).

**Figure 3.**
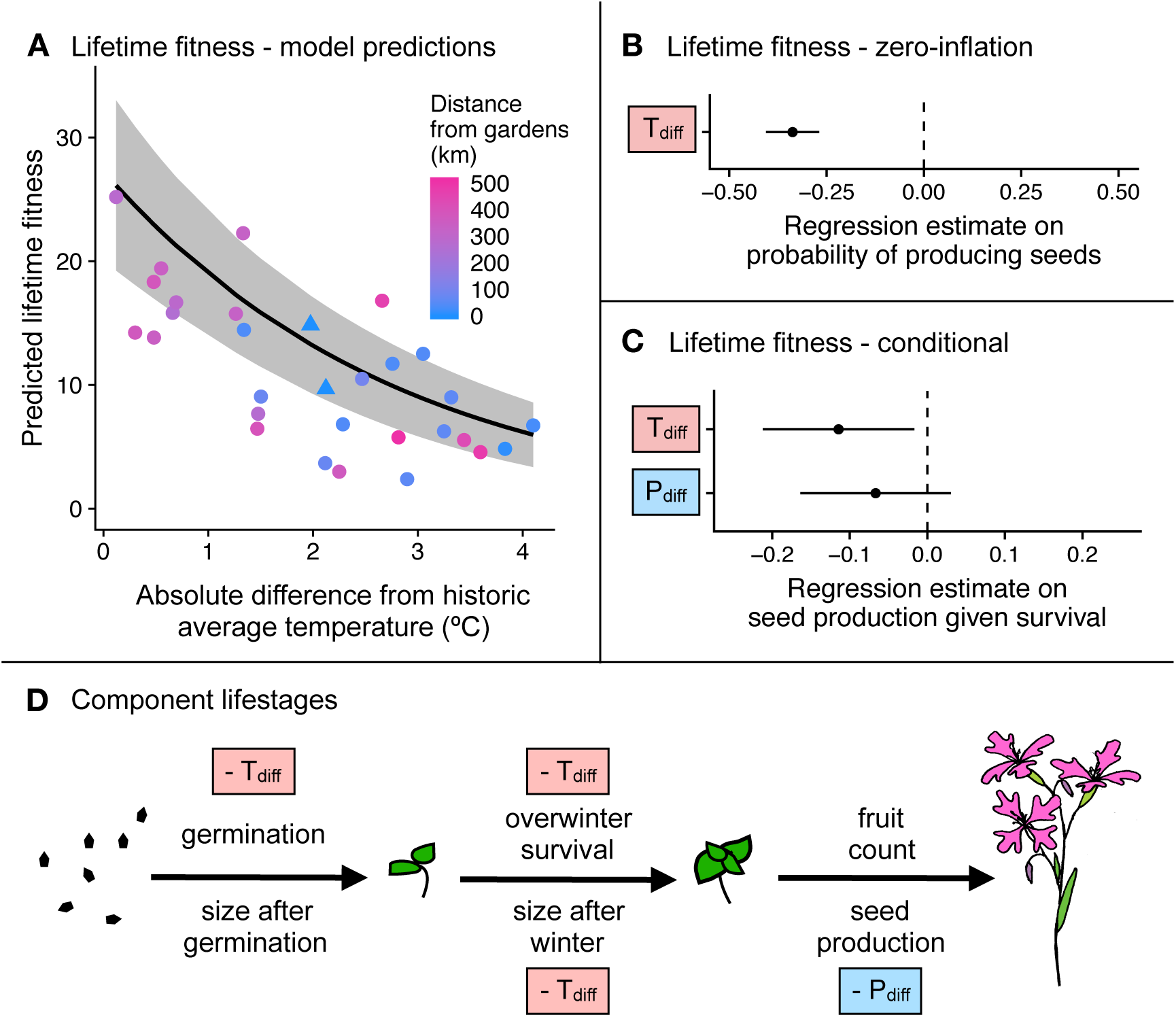
Effects of absolute temperature difference (September-July; T_diff_) and absolute precipitation difference (April-July; P_diff_) on performance of *Clarkia pulchella* in common gardens. These analyses include within-population plants only (no gene flow). **(A)** Lifetime fitness (number of seeds produced by each seed that was planted) declines with increasing differences in temperature between the historic average of the source population and the experimental conditions in the transplant gardens. The regression line and 95% confidence interval incorporate both the conditional and zero-inflation model components; the confidence interval is conditioned on fixed effects only. Though these temperature differences are expressed as absolute, almost all populations were from sources that are historically cooler than the transplant sites were during the experiment. Points are raw averages for each source population in each garden, colored by the distance between the source population and the transplant garden. Triangles are focal populations in their home sites. **(B)** Regression estimates and standard errors from the zero-inflation part of a model of lifetime fitness. **(C)** Regression estimates and standard errors from the conditional part of a model of lifetime fitness. **(D)** Schematic of effects of absolute temperature differences (T_diff_) and absolute precipitation difference (P_diff_) on component lifestages of *Clarkia pulchella*. Directionality of effects is illustrated with “-”; in these analyses all significant effects were negative. Predictors in boxes are significant (*P* < 0.05). Size in the previous lifestage is not shown here, but has a significant positive effect on overwinter and reproductive lifestages. This summarizes the significant results of separate models for each lifestage; full statistical results of these tests are in Table S3.

Analyses of component lifestages support these inferences (Figure 3D, Table S3): being poorly matched to experimental temperatures had negative effects on germination proportion, overwinter survival, and the size of plants after winter. While precipitation differences were not significant in the model of lifetime fitness (*β* = −0.067, SE = 0.049, *P* = 0.177, Figure 3C), they did have a negative effect on seed production among plants surviving the winter (Figure 3D, Table S3).

### Gene flow may confer some benefits to edge populations

As in the analyses of within-population plants only, both midparent temperature differences and midparent precipitation differences had negative effects on lifetime fitness in our common gardens (Table S4A; Figure 4AB). Gene flow (i.e., being a between-population rather than a within-population cross) did not have a significant effect in the lifetime fitness model that also included climate differences.

**Figure 4.**
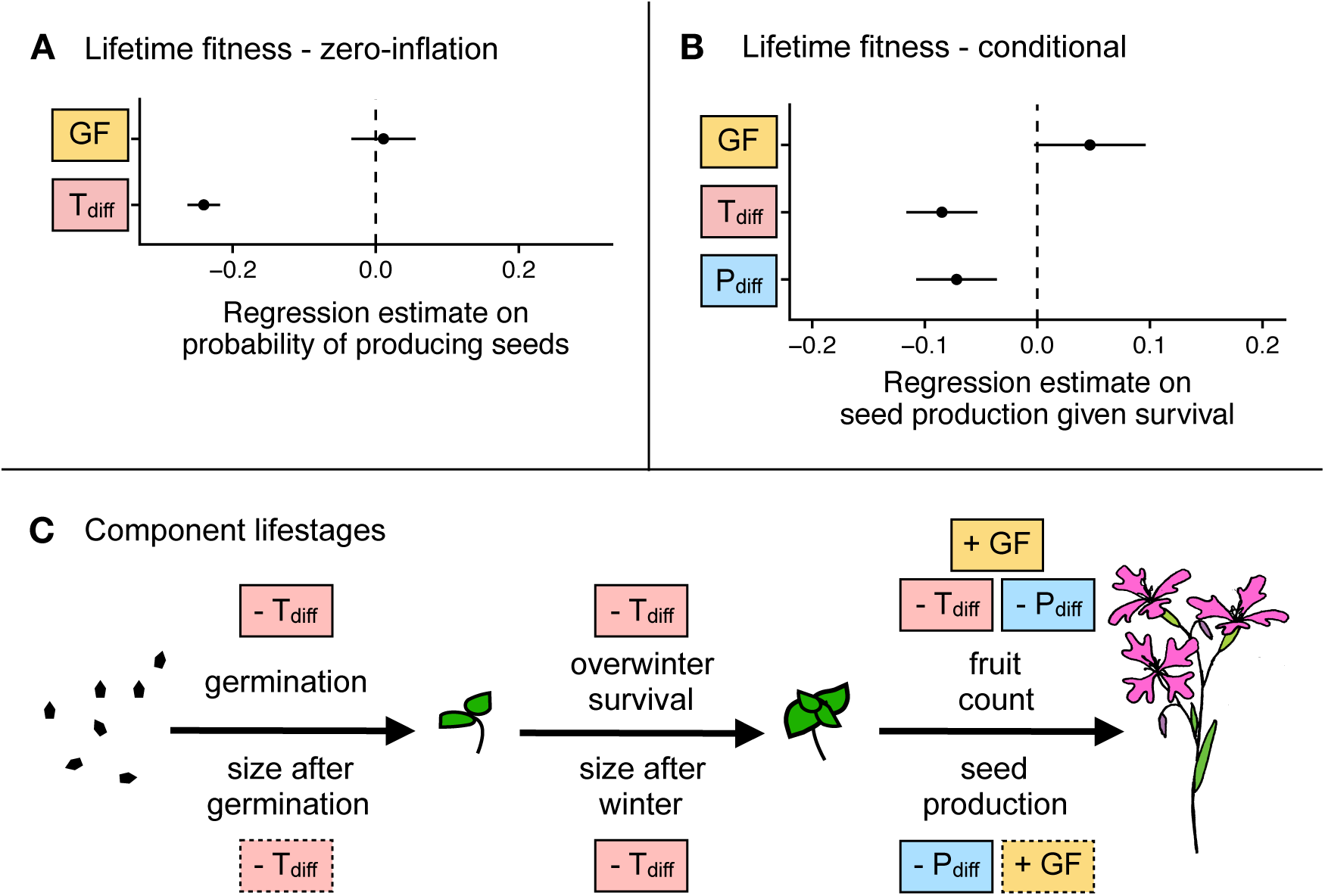
Effects of gene flow (differences between between-population and within-population crosses) on performance of *Clarkia pulchella*, accounting for midparent temperature and precipitation. Lifetime fitness (the number of seeds produced by each seed that was planted) incorporates germination, survival, and reproduction. **(A)** Regression estimates and standard errors from the zero-inflation part of a model of lifetime fitness. **(B)** Regression estimates and standard errors from the conditional part of a model of lifetime fitness. **(C)** Effects of gene flow (GF), absolute midparent temperature differences (T_diff_), and absolute midparent precipitation differences (P_diff_) on component lifestages of *Clarkia pulchella*. Directionality of effects is illustrated with “+” and “-”. Not quite significant parameters (0.05 < *P* < 0.10) are shown in boxes with dashed margins, predictors in solid boxes are significant (*P* < 0.05). Size in the previous lifestage is not shown here, but has a significant positive effect on overwinter and reproductive lifestages. This summarizes the significant results of separate models for each lifestage; full statistical results of these tests are in Table S4.

It is difficult to disentangle the effects of precipitation differences from the effects of gene flow in these analyses. This is because our focal populations are already among the driest provenances in our experiment. Therefore, the average between-population plant is better matched to the experimental conditions than the average within-population plant, because the midparent precipitation of between-population plants is always calculated with at least one very dry focal parent (Figure S3A). This was not an issue with temperature differences, because our focal populations are intermediate to other provenances in terms of temperature (Figure S3B). When lifetime fitness was analyzed without precipitation differences in the model, we found that gene flow (being a between-population cross, rather than a within-population cross), had a positive effect on lifetime fitness in addition to effects of temperature (Table S4B).

A small positive effect of gene flow, independent of climatic differences, is supported by analyses of some lifestage components (Figure 4C; Table S4C). Negative effects of precipitation and temperature differences were similar to those found in the analyses of climatic drivers of performance, while gene flow (i.e., being from a between-population vs. a within-population cross) had a positive effect on fruit production and a marginal positive effect on seed production.

### Genetic differentiation between parental populations positively affects fitness

Both midparent temperature difference from the garden conditions and genetic differentiation between parental populations had significant effects on fitness (Figure 5, Table S5AB, Table S6A). Genetic differentiation between parental populations had a positive effect on lifetime fitness via increasing the probability of producing seeds (the zero-inflation part of the model). The effects of genetic differentiation on lifetime fitness were mirrored in the analyses of single lifestages: F_ST_ had a positive effect on germination, size after winter, fruit count, and seed production (Figure 5C, Table S5A, Table S6A).

**Figure 5.**
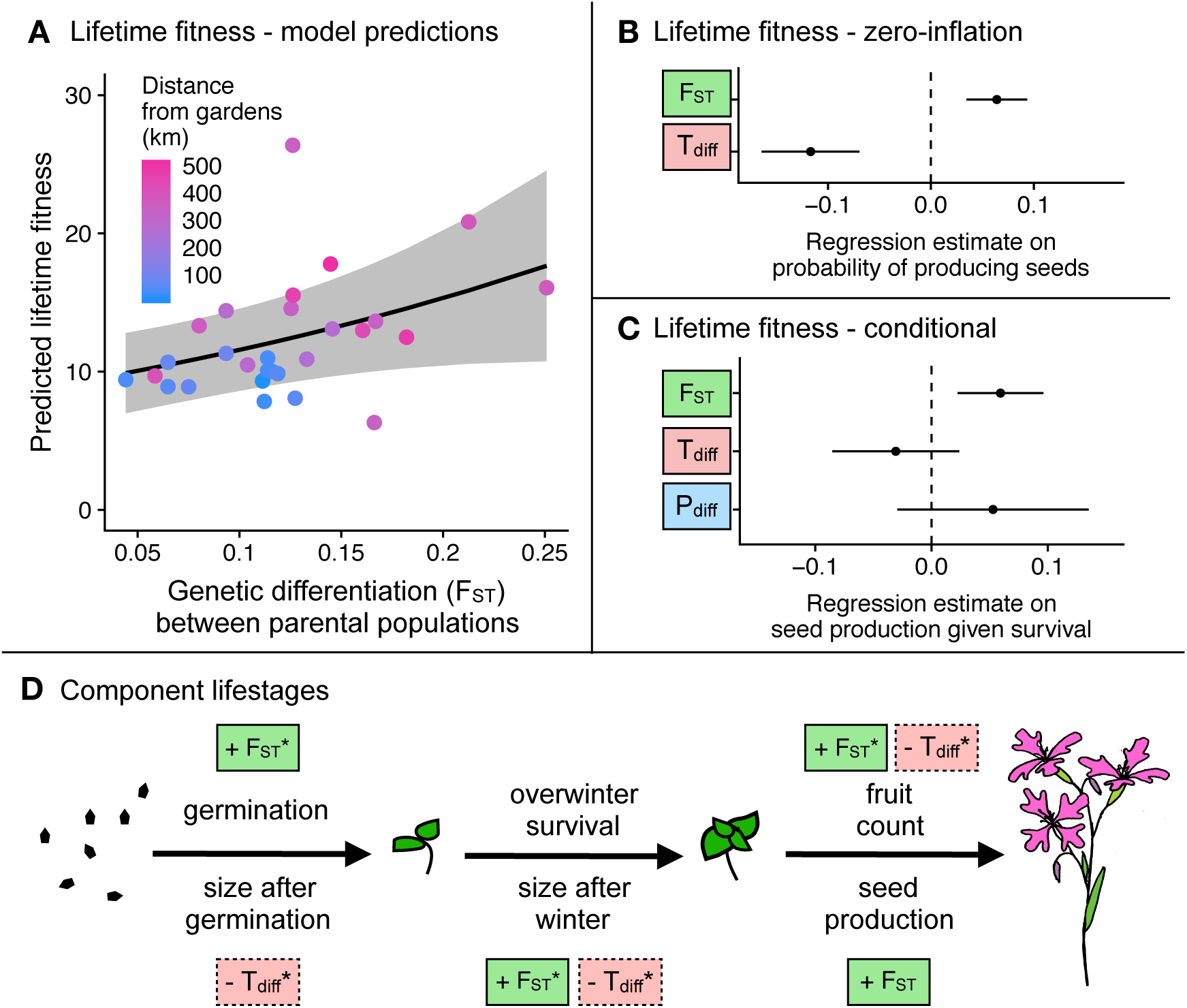
Effects of genetic differentiation between parental populations, as well as midparent temperature and precipitation on performance of *Clarkia pulchella*. **(A)** Among between-population crosses, increased genetic divergence between parental populations had a positive effect on lifetime fitness (number of seeds produced by each seed that was planted). The regression line and 95% confidence interval show the effects of F_ST_ when T_diff_ and P_diff_ are held constant. These incorporate both the conditional and zero-inflation model components; the confidence interval is conditioned on fixed effects only. Points are raw averages for each gene flow source in each garden, colored by the distance between the source population and the garden (the home of the recipient population). **(B)** Regression estimates and standard errors of genetic differentiation (F_ST_) and absolute midparent temperature differences (T_diff_) on the probability of producing seeds. **(C)** Effects of genetic differentiation, temperature differences, and absolute precipitation differences (P_diff_) on conditional seed production. **(D)** Effects of T_diff_ and F_ST_ on component lifestages of *Clarkia pulchella*. Precipitation differences were not significant when tested for component lifestages. Directionality of effects is illustrated with “+” and “-”. Not quite significant parameters (0.05 < *P* < 0.10) are shown in boxes with dashed margins, predictors in solid boxes are significant (*P* < 0.05). Size in the previous lifestage is not shown here, but has a significant positive effect on overwinter and reproductive lifestages. * indicates predictors that are only significant in separate models, not in full models with all predictors. Complete statistical results of these tests are in Table S5 and Table S6.

The negative effect of midparent temperature differences in between-population crosses is generally consistent with our analyses of climatic drivers of performance in within-population crosses. Between-population plants with donor parents that were well-matched to temperatures during the experiment were more likely to produce seeds, as indicated by the significant negative effect of midparent temperature differences in the zero-inflation part of that model (Table S5B, Table S6A).

Midparent temperature differences did not significantly affect any single lifestage, but had marginal negative effects on size after germination, size after winter, and fruit number (Figure 5C, Table S5B). Midparent precipitation differences did not significantly affect lifetime fitness or component lifestages (Table S5C).

It is important to note that genetic differentiation is not especially correlated with temperature differences (r = −0.25, *P* = 0.193), though genetic differentiation and precipitation differences are correlated (r = 0.64, *P* = 0.0003), with plants whose parents are more genetically differentiated also having larger differences between their historic midparent precipitation and conditions during the experiment. We think it is unlikely that the apparent positive effects of genetic differentiation are actually driven by precipitation differences, because we would expect high precipitation differences to negatively affect fitness. The overall weak or absent effects of temperature and precipitation differences in these models may be due to a narrower range of variation in midparent climate for the between-population crosses relative to the within-population crosses (Figure S3).

## Discussion

We conducted a common garden experiment at the northern range margin of *Clarkia pulchella* to examine how the effects of gene flow on peripheral populations vary with climatic and genetic differentiation between focal and source populations. We examined predictors of fitness of within-population crosses, in which both parents originated from the same source population, as well as between-population crosses, in which one parent was local to the common gardens and the other was from another population from across the northern half of the range of *C. pulchella*. In our experiment, provenances of *C. pulchella* from climates that most closely matched the conditions during the experiment performed best. Populations of *C. pulchella* at the northern range margin benefited from gene flow from warm source locations during the warm year of our experiment. Gene flow also conferred some benefits independent of climate, as evidenced by the positive effect of gene flow on later lifestages when controlling for temperature differences and the positive effect of increasing genetic differentiation between the parental populations.

## Climate of origin predicts performance

Populations of *Clarkia pulchella* are adapted to their historic climate regimes, a pattern consistent with findings in many other species (Anderson et al., 2015; Wilczek et al., 2014). When grown in common sites, the performance of individuals was determined by the degree to which conditions during the experiment deviated from historic temperature and precipitation averages of each provenance (Figure 3). Because of this local adaptation to climate, gene flow from sites that deviate from local conditions (in our experiment, sites that are cooler than the focal populations) had the potential to disrupt local adaptation, as indicated by the somewhat negative effects of midparent temperature differences on between-population plants (Figure 5, Table S5). Although simulated gene flow from divergent environments had the potential to reduce fitness in this experiment, it is unlikely that gene flow occurs at a high enough rate among natural populations to swamp local adaptation. However, our results highlight that gene flow and dispersal need not be from populations that are geographically distant (or from the center of the range) to be climatically divergent from historic or current conditions. Rather, two of the most temperature-mismatched populations used in the experiment are from sites nearest to our common gardens (populations D1 and D3; Figure 1).

Under climate change, local adaptation to historic climate regimes may generate local maladaptation in field trials. We see this in our results: populations from warmer locations performed best in our gardens (Figure 3, Table S3) and outperformed local populations (Supplementary Analysis 1 and Figure S5), and gene flow from warmer locations had positive effects on some lifestages (Figure 5, Table S5). This lagging adaptation to climate has been documented in other recent common garden studies. In a reciprocal transplant experiment of a long-lived sedge, McGraw et al. (2015) found that populations were displaced 140 km south of their optimal climate conditions. Wilczek et al. (2014) found that local genotypes of *Arabidopsis thaliana* from across Europe were consistently outperformed in common gardens by accessions from historically warmer locations. These results indicate that dispersal and gene flow are important processes promoting range stasis as climate warms, as they allow alleles that are beneficial in warm environments to spread from historically warm populations to recently warming sites. However, climate is multivariate, and as the climate changes it may generate combinations of conditions that no population has historically experienced (Williams and Jackson, 2007; Mahony et al., 2017). The particular combination of hot and dry conditions in our common gardens was unlike any of our populations’ historic temperature and precipitation combinations and approached future climate projections for these sites (Figure 1), though they are similar to normals from some populations not included in our experiment, primarily from the southern half of the species range (data not shown). While precipitation conditions were similar to those historically experienced by the focal populations, temperature conditions favored another set of populations. Whether the optimal traits for different climatic axes are antagonistic and whether segregation and recombination will allow adaptation to novel climates are important considerations in predicting climate change responses.

## Gene flow confers benefits independent of climate

We saw some additional positive effects of gene flow once the effects of climate are controlled for (Figure 4, Table S4). These positive effects may be the result of reduced homozygosity when parental plants come from two different populations; this inference is supported by the positive effect of genetic differentiation between parental populations on performance (Figure 5, Table S5). These results are also generally consistent with previous work in which experimental populations of *Clarkia pulchella* with higher genetic effective population sizes had lower extinction probabilities (Newman and Pilson, 1997). An interesting direction for future models of swamping gene flow along environmental gradients might be to explore whether incorporating heterosis-dependent increases in the effective migration rate (Ingvarsson and Whitlock, 2000) alters predictions (this could be done with various dispersal distances, under scenarios of various magnitudes of isolation-by-distance). However, an important question is whether the benefits of reduced homozygosity (or increased heterozygosity) are transient effects among F1s, how long they would persist in future generations if our between-population plants backcrossed into the focal populations, and whether these benefits may be counteracted by outbreeding depression as recombination disrupts co-adapted gene complexes. The answers to these questions are likely to depend on many factors, including the genetic architecture of local adaptation and population size (Willi et al., 2007), but fitness de-clines in subsequent generations are not uncommon after between-population crosses (Fenster and Galloway, 2000; Johansen-Morris and Latta, 2006). Novel environments may alter the costs and benefits of outbreeding: increases in variation among individuals might help populations adapt, despite temporary decreases in mean fitness due to outbreeding depression.

During this study, the effects of being well-matched to the experimental conditions seemed to dominate over potential benefits of being from a local population (for example, the benefits of being adapted to local soil conditions or herbivores). This inference is supported by the fact that lifetime fitness of local populations did not differ from that of foreign populations even once climate differences were controlled for (Supplementary Analysis 2, Table S7) though our experiment was not especially well-suited to test this because we have only two local populations.

## Limited inference about population persistence

Our ability to make inferences from our results about the longer-term effects of gene flow on the persistence and adaptive potential of range edge populations is limited. While it seems clear that gene flow from warm sites is likely to accelerate adaptation to warming conditions, we do not know whether these populations were historically limited by adaptation, and whether the additional genetic variation introduced by gene flow would permit better adaptation to local conditions and range expansion on an evolutionary time scale. These types of questions are difficult to test in field systems (but see Etterson and Shaw 2001), but inferences can be made by examining genetic variance of wild populations in the lab (Kellermann et al., 2006; Hoffmann et al., 2003). The development of experimental evolution systems to test equilibrial range dynamics is an exciting avenue for future work—this would be a natural extension of recent studies of range expansion dynamics using experimental evolution in the lab (Ochocki and Miller, 2017; Williams et al., 2016). It is also important to note that all populations in our experiment had reproductive rates that were well above replacement (one seed produced per seed planted, see y-axis on Figure 3A, Figure 5A), so we have no evidence that gene flow has the potential to drive populations extinct, or to rescue them from extinction. The high lifetime fitness we observed during our experiment could be due to several factors. First, perhaps warm conditions over the entire season are favorable for all sources, but are more favorable for warm-adapted populations. Alternatively, we could have increased fitness by limiting antagonistic biotic interactions. Additionally, our conversion from ovary length to seed production estimates assumes that pollinator service is at least as effective as our hand pollinations and that seed production is not resource-limited. If preventing pollinator access increased fruit production due to within-plant resource reallocation, our estimates of fitness from ovary length could be overestimates. Finally, and perhaps most plausibly, our plot placement may have upwardly biased our germination and reproductive estimates. We placed plots in patches that appeared favorable to *C. pulchella*, but naturally dispersing seeds are likely to land in a mix of favorable and unfavorable patches. We do not know whether any of these factors might interact with provenance, in which case they might change the relative performances of populations in our experiment.

## Conclusions and future directions

On heterogeneous landscapes, the effects of gene flow are likely to depend strongly on climate of origin, which may be decoupled from geography. Although gene flow from divergent climates may result in reduced fitness, gene flow can also confer benefits to range edge populations via both pre-adapted alleles and reduced homozygosity. As a result, moderate rates of gene flow likely represent a net benefit to edge populations, particularly as the environment is changing. This study highlights the challenges of testing hypotheses about equilibrial range limits in the field, where climate change is a persistent reality. Even if populations were once locally adapted, they are likely no longer at equilibrium with climate. The signal of climate anomalies disrupting local adaptation can be detected in published literature to date (Bontrager et al., in prep.). In light of this, future studies of local adaptation at range edges should be designed in such a way that the results will be informative even in non-equilibrial conditions.

## Acknowledgements

We would like to thank M. Osmond, L. Bontrager, C. Leven, D. Gamble, A. Porter, A. Wilkinson, J. Chan, T. Mitchell, and P. Chen for their help in the field. A. Wilkinson and M. Zink Yi provided plant care in the greenhouse. B. Gass and D. Holtz also helped with plant care. E. Fitz and D. Gamble assisted with greenhouse pollinations and fruit collection. A. Porter, J. Ono, and J. Chan assisted with preparing the seeds for planting. British Columbia Provincial Parks permitted the garden installation and monitoring. M. Whitlock, J. Whitton, S. Aitken and T. Usui provided helpful comments on an earlier draft of this manuscript. This work was supported by grants from the Botanical Society of America and the Washington Native Plant Society. MB was supported by a University of British Columbia Four-year Fellowship. The authors declare no conflicts of interest.

## Author contributions

MB and ALA conceived of the project. MB performed greenhouse crosses, garden installation, and data collection. MB analyzed the data in consultation with ALA. MB wrote the manuscript with advice and edits from ALA.

## Data deposition

All data and code required to recreate these analyses is publicly available at github.com/megan bontrager/clarkia-pulchella-transplant. Upon publication, all data and code will be archived in an appropriate repository (e.g., Dryad).

## Supplementary methods

### 1. Greenhouse conditions and crossing design

Seeds were planted in the greenhouse 9-11 December, 2014 in conetainers (Stuewe and Sons, Tangent, Oregon, USA) filled with Sunshine Mix No. 4 (Sun Gro Horticulture, Agawam, MA, USA). For each of 22 maternal families per population, 3-5 seeds were planted on the soil surface in each pot. For families from each of the two focal populations, three replicate pots were planted per family because larger quantities of flowers would be needed from these families; other populations were represented by one replicate per family. Pots were arranged into randomized blocks, with each block containing one family from each population (one pot from each donor population and three replicate pots from each of the two focal populations). The soil was kept moist until germination, then plants were hand watered every 1-3 days as needed to prevent wilting. After germination, plants were thinned randomly to one per cone and pumice was added to the soil surface to prevent fungal growth. Plants began to flower in March 2015. Plants were bagged to prevent unintentional pollination, and flowers were emasculated upon opening to prevent self-pollination. For the crosses, 20 of the 22 blocks were used, the other two were maintained in the same growing conditions to provide alternate plants in case of mortality or sterility. We performed as many crosses as possible using a single focal plant, but if flower production was too low on that plant we also used one of the replicate focal plants from the same family. Most crosses had to be performed 2-3 times to obtain adequate numbers of seeds for the experiment. Some crosses could not be performed due to mortality, sterility, or limited flower production. As ripening progressed, the ends of fruits were taped shut to prevent seed loss. Upon ripening, fruits were collected and stored in coin envelopes in the lab. Crosses were performed March-May 2015 and and we collected fruits March-June 2015.

### 2. Transplant installation

Seeds attached to toothpicks were planted in the ground 18-21 September 2015. Plots were prepared by removing litter, large rocks, and dried remains of herbaceous perennial plants. The ground surface was minimally leveled to allow for placement of planting grids that aided in consistently spacing the plants. Each toothpick was inserted into the ground gently so that seeds were not dislodged or damaged until seeds were ~3 mm below the soil surface. Toothpicks were inserted at 5 cm spacing into ~1 m by 2 m blocks. Block shape was varied to accommodate rocks and shrubs surrounding the planting area. After planting, each block was protected with 20 cm high hardware cloth cages supported by rebar. These cages were intended to prevent trampling by larger animals but did not prevent entry of rodents and other small animals. The area surrounding the plots at each site was sprayed with deer repellent several times during the course of the experiment. To ensure germination, plots were watered at a rate of ~10 L per plot 27-29 October 2015, though at that time most seeds that were checked already had radicles emerging. In May 2016 cattle fencing was put around the plots at the Blue Lake site before cattle were released into the area for grazing; this fencing succeeded in keeping the cows off the plots. No cattle were present at the Rock Creek site.

### 3. Details of monitoring and measuring

Our transplant gardens were installed in areas where *C. pulchella* occurs naturally, and because of this we wanted to evaluate how frequently we might have mistaken naturally occurring plants for experimental plants. During germination surveys we censused one more germinant than the number of seeds that we planted at 23 out of 16,680 grid points (0.14%). This gives an estimate of the minimum rate at which naturally occurring seeds were indistinguishable from our planted seedlings. So, while it is probable that some naturally occurring plants were mistaken for experimental plants, we consider the frequency of possible misidentification to be acceptably low.

During winter, some plots were affected by frost-heave and seedlings were uprooted from their planting locations when their toothpick was forced out of the ground (1901/16680 grid points, 11.4%). In lightly affected areas, toothpicks and seedlings were gently settled back into the soil. In more heavily affected areas, individual identity could no longer be determined confidently and individuals were excluded from further measurements and analyses (95/16680 grid points, 0.57%).

Censuses of reproduction began on 2 June 2016. On 12-13 June 2016 we censused spring survival of all plants. Once flowering began, we placed bridal-veil nets over the hardware cloth on each plot to prevent pollen escape into local populations. In June we censused each plot every 2-3 days. We recorded the date of first flowering of each plant at this temporal resolution. During each census, the immature ovary length of each new flower was recorded to be used as a proxy for maximum seed set. Flowers were marked as they were measured with a permanent marker and a running flower count was kept for each plant to avoid double-counting as flowers senesced. We continued these assessments as flowering slowed in July, but reduced the census interval to once a week. Damage to plants, such as rodent activity or herbivory, was noted during monitoring. Any plants with uncertain identities (due to frost damage as mentioned above, being far from their toothpick, or the toothpick disappearing; n = 201/16680 toothpicks, 1.2%) were excluded from all analyses. Plants that were killed by gophers, browsers, or galling insects were excluded from analyses that involved lifestages downstream from these events (n = 525/16479 plants, 3.2%) because we do not think that this mortality is related to population origin but rather to block-specific factors.

### 4. Ovary length as a fitness proxy

Pollinations were performed on a subset of plants to calibrate a conversion from immature ovary lengths to seed production. On 596 flowers (mean = 29.8 per plot, range = 0–126) stigmas were dusted with an ample pollen load using all four anthers from another plant in the plot. These flowers were marked with strings around the pedicles and fruits were collected when ripe. Seeds in each of these pollinated fruits were later counted in the lab. Total seed production per individual was estimated by multiplying the total ovary length of each plant by the average number of seeds per mm of immature ovary, as determined from the pollinated fruits. This resulted in an estimate of 4.75 seeds per mm based on a linear regression of number of seeds predicted by ovary length with the intercept set to 0 (Figure S6, 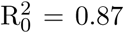, *P* < 0.0001, n = 596). This may be an over-or underestimate if our hand pollinations are more effective or less effective than natural pollinator service. For comparison, we measured ovary length and later seed set on a small number of naturally occurring *C. pulchella* plants near our plots (n = 73); the relationship between ovary length and number of seeds is similar in hand and natural pollinations (Figure S6).

We pollinated only a maximum of one flower per plant, so these may be overestimates because they do not account for potential resource limitation of seed set. If plants are prevented from setting seed due to pollen limitation, they may produce more flowers as a response. We found this to be the case to a small extent in natural populations of *Clarkia pulchella*, where pollinator exclusion led to plants producing an average of 0.6 more fruits compared to controls (Bontrager et al., 2018). However, we checked whether variation in seeds per mm of fruit in our transplanted individuals was associated with individual fitness (the overall fruit production per individual) or block quality (which we estimated based on the average fruit production of a block), and we could not attribute variation in seeds per mm of fruit to either of these factors. Therefore, while our conversion from fruit length to seeds may not be exact, we do not expect it to be systematically biased by plant size or resource availability.

An ANOVA indicated a significant effect of source population on seeds per mm of fruit for within-population plants (*P* = 0.040, F_14,298_ = 1.79) and a nearly significant effect for between-population plants (*P* = 0.060, F_14,298_ = 1.68). However, posthoc comparisons (Tukey HSD) did not reveal any significant pairwise differences. So while it is possible that there is interesting variation among populations in seed size or ovary morphology, we were not confident that we would be able to characterize this variation in sources where our hand pollination sample size is low, so we elected to use the same conversion across all populations.

We also tested whether seeds per mm of fruit covaried with any of the predictor variables that we used in our analyses, such that our conversion might amplify or dampen climate-fitness or climate-gene flow relationships. We found a small but significant difference between within-population and between-population plants in seeds per mm of fruit (within-population plants produced 0.8 fewer seeds per mm, on average; *P* < 0.0001). As a result, we may have slightly underestimated the positive effects of gene flow on seed production. We also found a small but significant effect of historic precipitation of each source population on seeds per mm of fruit, with populations from the driest site in the experiment producing on average 2.14 more seeds per mm of fruit than those from the wettest site (*P* < 0.0001). This could reflect either adaptive differentiation in seed size in response to precipitation, or that populations that were best matched to the experiment conditions (those from dry sources) not only made larger fruits but also produced more seeds per mm of fruit. In this case, our use of ovary length as a proxy for seed set is conservative, as it might have dampened the fitness benefits of being well matched to precipitation, rather than inflating them. We found no effects of temperature of origin or F_ST_ on seeds produced per mm of fruit.

## Supplementary analyses and results

### 1. Do warmer foreign populations outperform local populations?

Our main analyses indicate that populations from historic climates better matched to the experimental conditions performed best. This raises the question of whether foreign populations from warmer sites outperform local populations when they are compared directly (as well as whether cooler foreign populations perform worse than local populations). To test this, we designated foreign populations as either warmer or cooler than local populations by comparing their historic temperature normals to the historic temperature normals of our focal populations. We then ran a zero-inflated negative binomial GLMM comparing the lifetime fitness of local populations, warmer foreign populations, and cooler foreign populations. We included random effects of site within block and sire population (models with a more complex random effect structure would not converge).

We found that warmer foreign populations had a higher probability of producing seeds than the local populations (zero-inflation component: *β* = 0.344, SE = 0.126, *P* = 0.006, Figure S5A; conditional component: *β* = 0.153, SE = 0.136, *P* = 0.260, Figure S5B). Similarly, cooler foreign populations had a lower probability of producing seeds than the local populations (zero-inflation component: *β* = −0.316, SE = 0.129, *P* = 0.015, Figure S5A; conditional component: *β* = 0.000, SE = 0.136, *P* = 0.999, Figure S5B). These results support the inference that temperature of origin is an important driver of performance, and show that this leads to categorical differences between local, warmer foreign, and cooler foreign populations, (Figure S5C).

### 2. Are there benefits of being local once climate of origin is controlled for?

If the focal populations in this experiment are locally adapted to conditions other than the climate variables that we have considered here, they may have fitness advantages over foreign populations once climate of origin has been accounted for. To test this, we re-ran our analyses of local vs. foreign performance and included absolute temperature and precipitation differences as covariates (essentially combining the first two analyses described in the main text methods). Our statistical methods were the same as in our main analyses: for lifetime fitness responses we used a GLMM with a zero-inflated negative binomial distribution. We also built individual models of component lifestages in which we used climate data from only the months in each census window. We did not find any significant differences between local and foreign fitness, even once the effects of climate were accounted for (Table S7).

## Supplementary tables and figures

**Table S1.** Geographic information for the populations of *Clarkia pulchella* used in this experiment.

| Population name | Map ID | Type | Latitude | Longitude | Elevation (m) |
| --- | --- | --- | --- | --- | --- |
| Blue Lake | F1 | Focal | 49.05 | -119.56 | 842 |
| Johnstone Creek | F2 | Focal | 49.04 | -119.05 | 866 |
| Border | D1 | Donor | 48.98 | -118.99 | 1211 |
| Day Creek Road | D2 | Donor | 48.94 | -118.51 | 911 |
| Bodie Mountain | D3 | Donor | 48.83 | -118.83 | 1603 |
| Boulder Creek Road | D4 | Donor | 48.76 | -118.33 | 1115 |
| Aeneas Valley | D5 | Donor | 48.54 | -118.91 | 1126 |
| Henry Creek Road | D6 | Donor | 47.45 | -114.77 | 1103 |
| Heyburn State Park | D7 | Donor | 47.34 | -116.79 | 801 |
| McCrosky State Park | D8 | Donor | 47.09 | -116.98 | 1186 |
| Graves Creek Road | D9 | Donor | 46.80 | -114.41 | 1201 |
| Bitterroot | D10 | Donor | 46.54 | -113.89 | 1424 |
| Abel's Ridge | D11 | Donor | 46.28 | -117.60 | 1457 |
| Tucannon | D12 | Donor | 46.24 | -117.74 | 1022 |
| Pendleton | D13 | Donor | 45.74 | -118.25 | 649 |

**Table S2.** Results of generalized linear mixed effects models for the effect of local vs. foreign origin on performance of *Clarkia pulchella* in common gardens. There are no significant differences between populations of local vs. foreign origin in fitness components or lifetime fitness. Size during the previous census (November for overwinter survival and size, March for fruit counts and estimated seed production) is always a significant predictor of performance in subsequent lifestages.

| Response | Foreign vs. local origin |  |  | Size during previous census |  |  |
| --- | --- | --- | --- | --- | --- | --- |
|  | Estimate | SE | *P*-value | Estimate | SE | *P*-value |
| Germination | -0.034 | 0.088 | 0.702 | - | - | - |
| Size after germination | 0.027 | 0.073 | 0.708 | - | - | - |
| Overwinter survival | 0.176 | 0.126 | 0.161 | **0.377** | **0.033** | **< 0.001** |
| Size after winter | -0.068 | 0.182 | 0.708 | **0.750** | **0.042** | **< 0.001** |
| Fruit count | 0.089 | 0.146 | 0.544 | **0.544** | **0.041** | **< 0.001** |
| Seed production | -0.005 | 0.128 | 0.966 | **0.388** | **0.032** | **< 0.001** |
| Lifetime fitness - zero inflation | -0.032 | 0.123 | 0.798 | - | - | - |
| Lifetime fitness - conditional | -0.093 | 0.128 | 0.469 | - | - | - |

**Table S3.** Results of generalized linear mixed effects models of the effects of absolute precipitation and temperature differences on component lifestages of *Clarkia pulchella*. Temperature and precipitation differences refer to absolute differences between the historic conditions that a population experienced and the conditions in the common gardens during the experiment. These differences were calculated using climate data from only the months of that census period (i.e., September-November for germination and size after germination, December-March for overwinter survival and size after winter, and April-July for fruit counts and seed production). Analyses were conducted using only plants surviving the previous census window. Whenever applicable, size in the previous census was included as a covariate to account for differences accumulated during earlier lifestages. Significant parameters are indicated with bold text. These results are visually summarized in Figure 3.

| Response | Absolute temperature difference |  |  | Absolute precipitation difference |  |  | Size in previous census |  |  |
| --- | --- | --- | --- | --- | --- | --- | --- | --- | --- |
|  | Estimate | SE | *P*-value | Estimate | SE | *P*-value | Estimate | SE | *P*-value |
| Germination | **-0.094** | **0.035** | **0.007** | - | - | - | - | - | - |
| Size after germination | -0.027 | 0.026 | 0.293 | - | - | - | - | - | - |
| Overwinter survival | **-0.078** | **0.032** | **0.015** | - | - | - | **0.366** | **0.033** | **< 0.001** |
| Size after winter | **-0.518** | **0.095** | **< 0.001** | - | - | - | **0.748** | **0.042** | **< 0.001** |
| Fruit count | -0.114 | 0.071 | 0.108 | -0.091 | 0.058 | 0.119 | **0.543** | **0.041** | **< 0.001** |
| Seed production | -0.055 | 0.047 | 0.249 | **-0.110** | **0.044** | **0.011** | **0.389** | **0.033** | **< 0.001** |

**Table S4.** Results of generalized linear mixed effects models of the effects of being a within-population cross vs. a between-population cross, while accounting for effects of absolute precipitation and temperature differences. Temperature and precipitation differences refer to absolute differences between the average historic conditions of an individual’s parental populations and the conditions in the common gardens during the experiment. Positive estimates of the effects of between-population vs. within-populations indicate that having parents from two different populations (“gene flow”) is beneficial. **(A)** Effects on lifetime fitness of *Clarkia pulchella*. **(B)** Effects on lifetime fitness when midparent precipitation differences are not included in the model. **(C)** Effects on component lifestages. Climate differences were calculated using climate data from only the months of that census period (i.e., September-November for germination and size after germination, December-March for overwinter survival and size after winter, and April-July for fruit counts and seed production). Analyses were conducted using only plants surviving the previous census window. Whenever applicable, size in the previous census was included as a covariate to account for differences accumulated during earlier lifestages. Significant parameters are indicated with bold text. These results are visually summarized in Figure 4.

| Response | Between-populations vs. within-populations |  |  | Absolute midparent temperature difference |  |  | Absolute midparent precipitation difference |  |  | Size during previous census |  |  |
| --- | --- | --- | --- | --- | --- | --- | --- | --- | --- | --- | --- | --- |
|  | Estimate | SE | *P*-value | Estimate | SE | *P*-value | Estimate | SE | *P*-value | Estimate | SE | *P*-value |
| **A. Lifetime fitness** |  |  |  |  |  |  |  |  |  |  |  |  |
| Lifetime fitness - zero inflation | 0.008 | 0.047 | 0.860 | **-0.240** | **0.023** | **< 0.001** | - | - | - | - | - | - |
| Lifetime fitness - conditional | 0.047 | 0.049 | 0.345 | **-0.085** | **0.032** | **0.007** | **-0.072** | **0.036** | **0.045** | - | - | - |
| **B. Lifetime fitness without precipitation in model** |  |  |  |  |  |  |  |  |  |  |  |  |
| Lifetime fitness - zero inflation | 0.008 | 0.047 | 0.860 | **-0.240** | **0.023** | **< 0.001** | - | - | - | - | - | - |
| Lifetime fitness - conditional | **0.114** | **0.036** | **0.002** | **-0.059** | **0.029** | **0.041** | - | - | - | - | - | - |
| **C. Component lifestages** |  |  |  |  |  |  |  |  |  |  |  |  |
| Germination | -0.035 | 0.078 | 0.653 | **-0.088** | **0.029** | **0.003** | - | - | - | - | - | - |
| Size after germination | -0.033 | 0.041 | 0.421 | -0.028 | 0.015 | 0.058 | - | - | - | - | - | - |
| Overwinter survival | 0.012 | 0.047 | 0.794 | **-0.058** | **0.022** | **0.009** | - | - | - | **0.336** | **0.023** | **< 0.001** |
| Size after winter | -0.052 | 0.127 | 0.678 | **-0.274** | **0.068** | **< 0.001** | - | - | - | **0.721** | **0.031** | **< 0.001** |
| Fruit count | **0.125** | **0.055** | **0.021** | **-0.081** | **0.032** | **0.011** | **-0.092** | **0.034** | **0.007** | **0.493** | **0.030** | **< 0.001** |
| Seed production | 0.092 | 0.047 | 0.052 | -0.030 | 0.028 | 0.285 | **-0.092** | **0.030** | **0.002** | **0.357** | **0.024** | **< 0.001** |

**Table S5.**
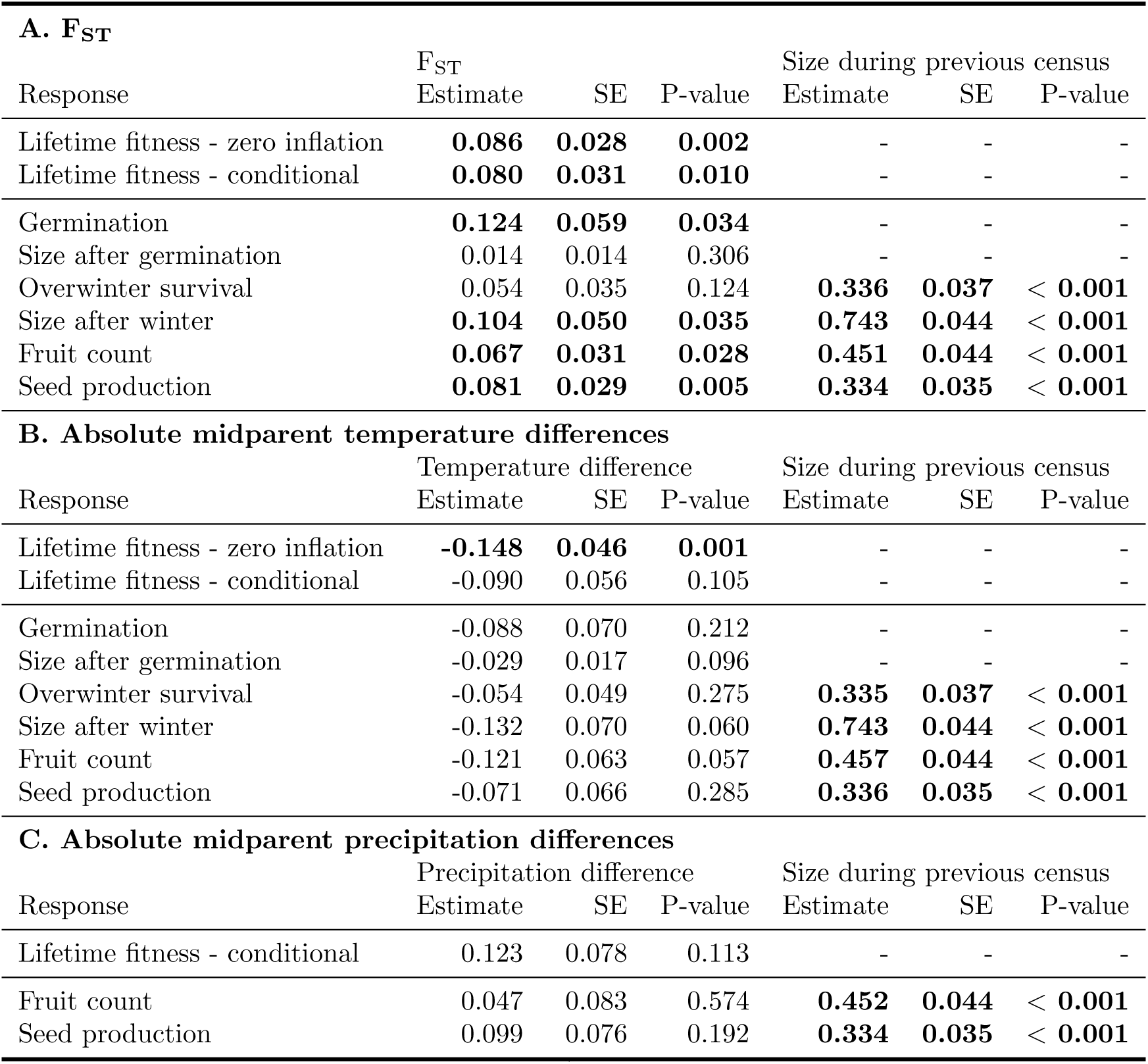
Results of generalized linear mixed effects models separately testing the effects of **(A)** genetic differentiation, **(B)** absolute midparent temperature differences, and **(C)** absolute midparent precipitation differences on performance of *Clarkia pulchella* in common gardens. Absolute midparent temperature and precipitation differences refer to absolute differences between the conditions in the common gardens during the experiment and the average historic conditions of an individual’s parental populations. These analyses were performed using between-population crosses only, that is, every plant has one parent from a focal population and one parent from a donor population. For analyses of lifetime fitness, temperature differences were calculated using the duration of the experiment and precipitation differences were calculated using summed April-July values. Precipitation differences are only included as an effect in the conditional part of the model of lifetime fitness because precipitation effects are expected to manifest at later lifestages. For analyses of component lifestages, climate differences were calculated using climate data from only the months of that census period (i.e., September-November for germination and size after germination, December-March for overwinter survival and size after winter, and April-July for fruit counts and seed production). Component lifestage analyses were conducted using only plants surviving the previous census window. Whenever applicable, size in the previous census was included as a covariate to account for differences accumulated during earlier lifestages. Significant parameters are indicated with bold text. These results are visually summarized in Figure 5.

**Table S6.** Results of generalized linear mixed effects models of the effects of genetic differentiation between parental populations on performance of *Clarkia pulchella* in common gardens. Effects of absolute precipitation and temperature differences are also included in these models. Temperature and precipitation differences refer to the absolute midparent differences, i.e., the absolute differences between the conditions in the common gardens during the experiment and the average historic conditions of an individual’s parental populations. These analyses were performed using between-population crosses only, that is, every plant has one parent from a focal population and one parent from a donor population. **(A)** Effects on lifetime fitness. **(B)** Effects on component lifestages. Climate differences were calculated using climate data from only the months of that census period (i.e., September-November for germination and size after germination, December-March for overwinter survival and size after winter, and April-July for fruit counts and seed production). Analyses were conducted using only plants surviving the previous census window. Whenever applicable, size in the previous census was included as a covariate to account for differences accumulated during earlier lifestages. Significant parameters are indicated with bold text. These results are visually summarized in Figure 5.

| Response | F<sub>ST</sub> |  |  | Temperature difference |  |  | Precipitation difference |  |  | Size during previous census |  |  |
| --- | --- | --- | --- | --- | --- | --- | --- | --- | --- | --- | --- | --- |
|  | Estimate | SE | P-value | Estimate | SE | P-value | Estimate | SE | P-value | Estimate | SE | P-value |
| **A. Lifetime fitness** |  |  |  |  |  |  |  |  |  |  |  |  |
| Lifetime fitness - zero inflation | **0.064** | **0.030** | **0.030** | **-0.117** | **0.048** | **0.014** | - | - | - | - | - | - |
| Lifetime fitness - conditional | 0.059 | 0.037 | 0.108 | -0.031 | 0.055 | 0.574 | 0.053 | 0.082 | 0.520 | - | - | - |
| **B. Component lifestages** |  |  |  |  |  |  |  |  |  |  |  |  |
| Germination | 0.111 | 0.062 | 0.073 | -0.042 | 0.080 | 0.798 | - | - | - | - | - | - |
| Size after germination | 0.005 | 0.016 | 0.770 | -0.026 | 0.020 | 0.179 | - | - | - | - | - | - |
| Overwinter survival | 0.047 | 0.044 | 0.286 | -0.014 | 0.060 | 0.809 | - | - | - | **0.335** | **0.037** | **< 0.001** |
| Size after winter | 0.072 | 0.061 | 0.232 | -0.073 | 0.082 | 0.376 | - | - | - | **0.742** | **0.044** | **< 0.001** |
| Fruit count | 0.058 | 0.034 | 0.089 | -0.093 | 0.058 | 0.110 | -0.010 | 0.084 | 0.905 | **0.455** | **0.044** | **< 0.001** |
| Seed production | **0.071** | **0.033** | **0.031** | -0.035 | 0.055 | 0.522 | 0.028 | 0.077 | 0.719 | **0.334** | **0.035** | **< 0.001** |

**Table S7.** Results of generalized linear mixed effects models for the effect of local vs. foreign origin on performance of *Clarkia pulchella* including covariates of absolute precipitation and temperature differences (Supplementary Analysis 2). Temperature and precipitation differences refer to absolute differences between the historic conditions that a population experienced and the conditions in the common gardens during the experiment. Analyses of lifetime fitness **(A)** use temperature differences over the entire growing period (September-July) and precipitation differences during spring and summer (April-July). Analyses of component lifestages **(B)** use climate data from only the months of that census period (i.e., September-November for germination and size after germination, December-March for overwinter survival and size after winter, and April-July for fruit counts and seed production), and these analyses were conducted using only plants surviving the previous census window. Size during the previous census (November for overwinter survival and size, March for fruit counts and estimated seed production) was also included as a covariate to account for differences accumulated during earlier lifestages. Significant parameters are indicated with bold text.

| Response | Foreign vs. local origin |  |  | Temperature difference |  |  | Precipitation difference |  |  | Size during previous census |  |  |
| --- | --- | --- | --- | --- | --- | --- | --- | --- | --- | --- | --- | --- |
|  | Estimate | SE | P-value | Estimate | SE | P-value | Estimate | SE | P-value | Estimate | SE | P-value |
| **A. Lifetime fitness** |  |  |  |  |  |  |  |  |  |  |  |  |
| Lifetime fitness - zero inflation | 0.025 | 0.124 | 0.840 | **-0.342** | **0.033** | **< 0.001** | - | - | - | - | - | - |
| Lifetime fitness - conditional | -0.137 | 0.135 | 0.312 | **-0.122** | **0.053** | **0.021** | -0.085 | 0.054 | 0.112 | - | - | - |
| **B. Component lifestages** |  |  |  |  |  |  |  |  |  |  |  |  |
| Germination | -0.024 | 0.085 | 0.776 | **-0.100** | **0.034** | **0.003** | - | - | - | - | - | - |
| Size after germination | 0.026 | 0.073 | 0.721 | -0.027 | 0.026 | 0.302 | - | - | - | - | - | - |
| Overwinter survival | 0.229 | 0.127 | 0.072 | **-0.087** | **0.032** | **0.007** | - | - | - | **0.367** | **0.033** | **< 0.001** |
| Size after winter | -0.061 | 0.176 | 0.728 | **-0.515** | **0.095** | **< 0.001** | - | - | - | **0.748** | **0.042** | **< 0.001** |
| Fruit count | -0.009 | 0.155 | 0.954 | -0.112 | 0.070 | 0.110 | -0.099 | 0.063 | 0.115 | **0.539** | **0.041** | **< 0.001** |
| Seed production | -0.101 | 0.134 | 0.448 | -0.064 | 0.051 | 0.216 | **-0.123** | **0.048** | **0.010** | **0.388** | **0.033** | **< 0.001** |

**Figure S1.**
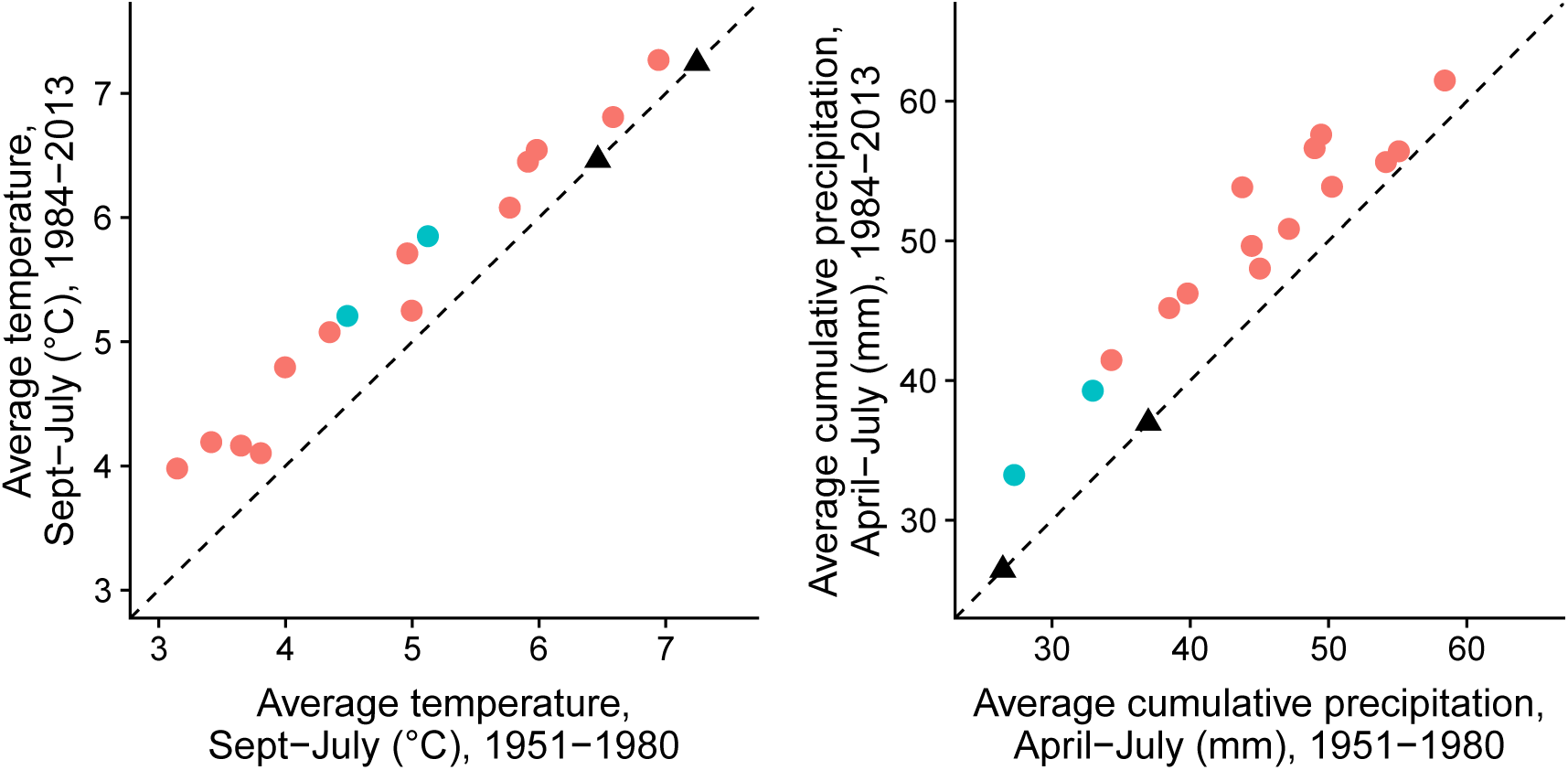
Correlation of temperature and precipitation normals calculated over the years 1951-1980 and 1984-2013. Each colored dot represents one source population, black dashed lines represent a 1:1 relationship, and black triangles represent conditions in the gardens during the experiment. Blue dots are focal populations, red dots are other populations. Climate data is from PRISM (PRISM Climate Group, 2017).

**Figure S2.**
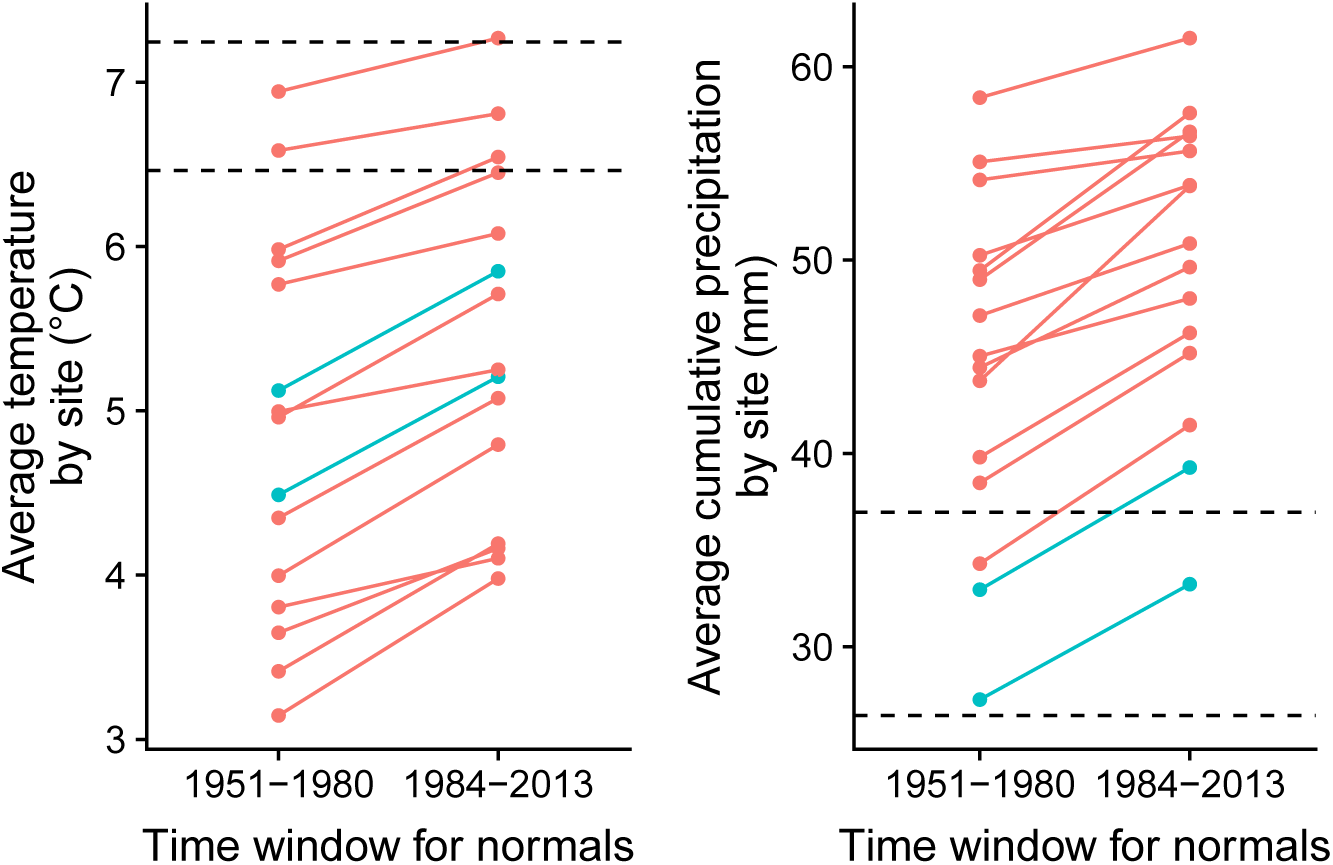
Temperature and precipitation normals calculated over the years 1951-1980 and 1984-2013, shown paired by site. Dashed black lines are the garden conditions. Blue dots are focal populations, red dots are other populations. Climate data is from PRISM (PRISM Climate Group, 2017).

**Figure S3.**
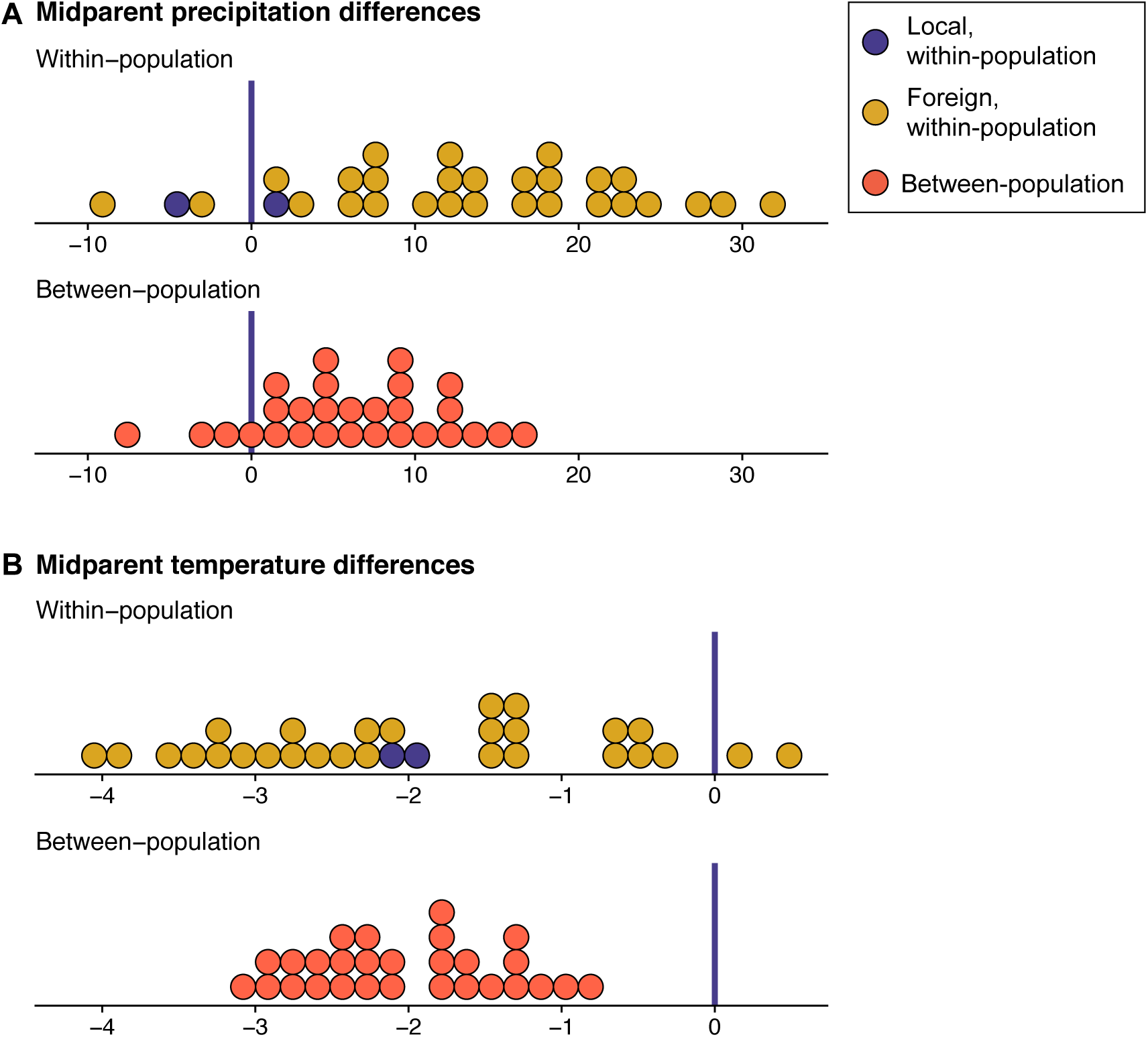
Distribution of climate differences of within-vs. between-population crosses of *Clarkia pulchella* relative to conditions during the experiment. Each dot represents a combination of maternal population, paternal population, and transplant site. Dark blue dots are within-population crosses of focal populations transplanted into their home sites. Gold dots are within-population crosses from donor populations planted into each of the two gardens, as well as the focal populations planted into each other’s sites. Red dots are between-population crosses. Vertical blue bars are placed at zero, indicating where populations would be perfectly matched to the temperature or precipitation condition as during the experiment. **(A)** Distribution of the differences between the average temperature in the home sites of parental populations and conditions during the experiment. Focal populations are intermediate in temperature relative to other populations in the experiment; this results in similar average differences in temperature in between-population crosses and within-population crosses. **(B)** Distribution of differences between the average precipitation in the home sites of parental populations and conditions during the experiment. Focal populations are among the driest in the experime e cnt; this results in smaller average differences in precipitation in between-population crosses compared to within-population crosses. Note that figures in the main text use absolute temperature differences: the absolute value of the midparent differences as they are plotted in this figure.

**Figure S4.**
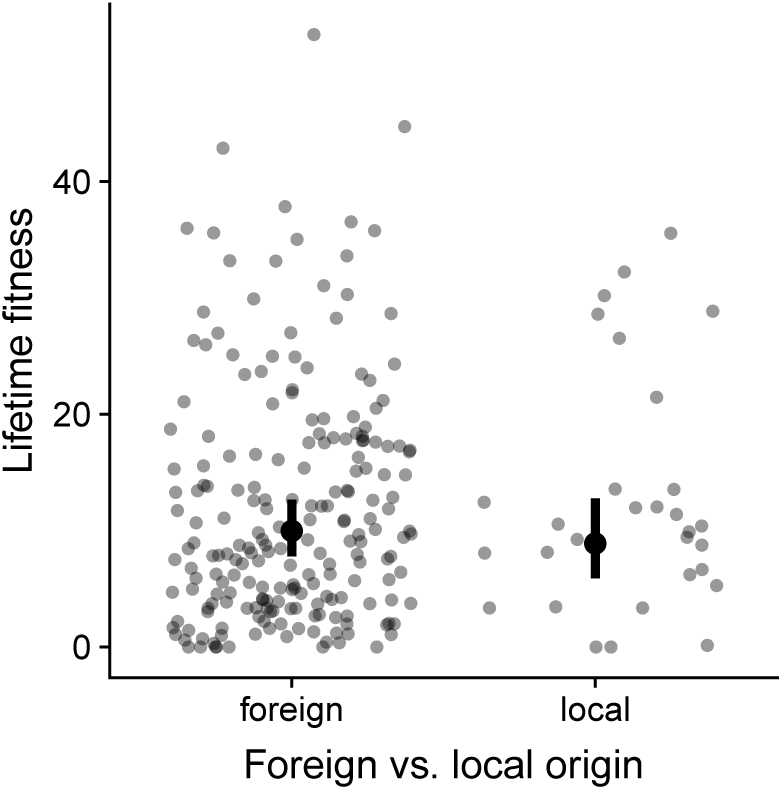
Lifetime fitness (seeds produced per seed planted) from populations of *Clarkia pulchella* with foreign vs. local parents. This analysis includes within-population plants only (no gene flow). Each point represents the average of a single family. Error bars are 95% confidence intervals of model estimated means, omitting variation from random effects.

**Figure S5.**
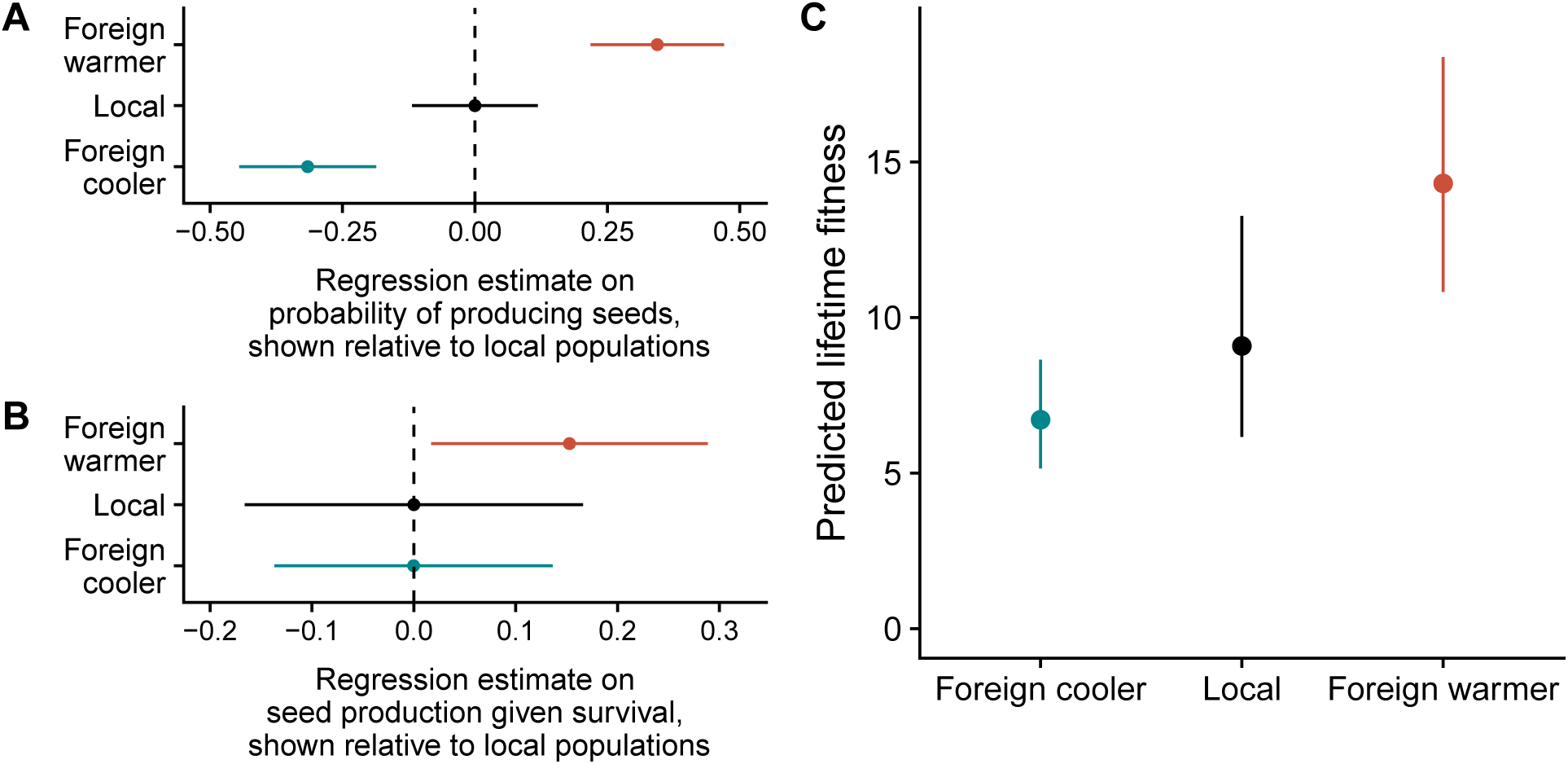
Regression estimates and predicted lifetime fitness of local populations (black) compared to foreign populations from warmer (red) and cooler (blue) provenances (Supplementary Analysis 1). **A** Estimated effects and standard errors of being from a warmer or cooler provenance on the probability of producing any seeds (the zero-inflation component of the model). **B** Estimated effects and standard errors of being from a warmer or cooler provenance on seed production, given survival (the conditional component of the model). Estimates in **A** and **B** are shown relative to local populations, which are set at 0. **C** Predicted lifetime fitness (seed produced per seed planted) of local and foreign populations from warmer or cooler provenances. These predictions integrate zero-inflation and conditional model components to give overall lifetime fitness projections (i.e. these predictions include seeds that did not survive to reproduce.)

**Figure S6.**
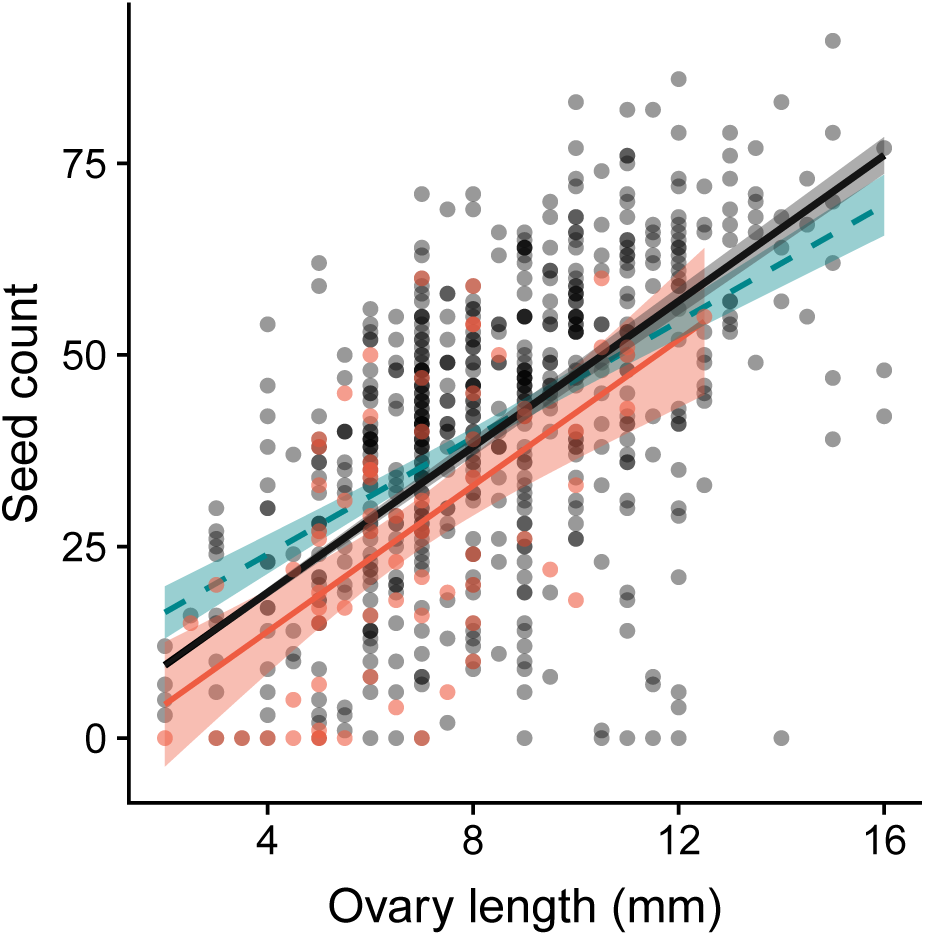
Immature ovary length (measured in the field) correlates with seed set after hand pollination (n = 596 fruits, grey points). This data was used to calibrate a conversion between ovary length in the field and potential seed production. The black regression line shows the relationship when the y-intercept is held at 0 (this line was used for our conversion); the dashed blue line shows the relationship when the intercept is not restricted. The red line and points show the relationship between ovary length and seed production in naturally occurring and naturally pollinated *Clarkia pulchella* in our common garden sites (n = 73 fruits).

